# The frequency and topology of pseudoorthologs

**DOI:** 10.1101/2021.02.17.431499

**Authors:** Megan L. Smith, Matthew W. Hahn

## Abstract

Phylogenetics has long relied on the use of orthologs, or genes related through speciation events, to infer species relationships. However, identifying orthologs is difficult because gene duplication can obscure relationships among genes. Researchers have been particularly concerned with the insidious effects of pseudoorthologs—duplicated genes that are mistaken for orthologs because they are present in a single copy in each sampled species. Because gene tree topologies of pseudoorthologs may differ from the species tree topology, they have often been invoked as the cause of counterintuitive results in phylogenetics. Despite these perceived problems, no previous work has calculated the probabilities of pseudoortholog topologies, or has been able to circumscribe the regions of parameter space in which pseudoorthologs are most likely to occur. Here, we introduce a model for calculating the probabilities and branch lengths of orthologs and pseudoorthologs, including concordant and discordant pseudoortholog topologies, on a rooted three-taxon species tree. We show that the probability of orthologs is high relative to the probability of pseudoorthologs across reasonable regions of parameter space. Furthermore, the probabilities of the two discordant topologies are equal and never exceed that of the concordant topology, generally being much lower. We describe the species tree topologies most prone to generating pseudoorthologs, finding that they are likely to present problems to phylogenetic inference irrespective of the presence of pseudoorthologs. Overall, our results suggest that pseudoorthologs are less of a problem for phylogenetics than currently believed, which should allow researchers to greatly increase the number of genes used in phylogenetic inference.

**Significance Statement:** Phylogenetics has long relied on the use of orthologs, or genes related through speciation events, to infer species relationships. However, filtering datasets to include only orthologs is both difficult and restrictive, drastically limiting the amount of data available for phylogenetic inference. Here, we introduce a model to study the probability and topologies of pseudoorthologs—duplicated genes that are mistaken for orthologs because they are present in a single copy in each sampled species. We show that pseudoorthologs are rare and that, even when they are present, they should not mislead phylogenetic inference. Our results suggest that strict filtering to remove pseudoorthologs unnecessarily limits the amount of data used in phylogenetic inference.

## Introduction

Phylogenetics aims to reconstruct evolutionary relationships among species. Recent advances in sequencing technologies have drastically increased the amount of data available for phylogenetic inference (1), which has led in turn to increased concern about how to filter large genomic and transcriptomic datasets. Central to most data-generating pipelines is the identification of orthologs, or genes related through speciation events, to the exclusion of paralogs, or genes related through duplication events (2). Because orthologous gene trees reflect only the species history, it has been argued that solely orthologs are appropriate for phylogenetic inference (e.g. 3, 4). Methods to extract orthologs from large datasets have therefore proliferated (reviewed in (5), e.g. (6–10)), but the task remains difficult, and pseudoorthologs (11) (or “hidden paralogs” (12)), are thought to represent a particularly insidious problem. Pseudoorthologs are paralogs that are mistaken as orthologs because, due to patterns of differential duplication and loss, they are present in a single copy in each sampled species.

Pseudoortholog gene trees can differ from the species tree in their topology and branch lengths. Consider, for example, a scenario in which a duplication occurred in the ancestor of three species (A, B, and C), where species A and B are sister species (Figure 1a,b). If one of the two copies is lost immediately, we can only sample genes with orthologous relationships (Figure 1c). If one copy is retained in species A and species B, while the other is retained in species C, then we have a pseudoortholog that is topologically identical to the true ortholog, but which has a longer internal branch (Figure 1d). Finally, if one copy is retained in species A (or B) and the other is retained in species B (or A) and C, then we have a pseudoortholog with a topology that differs from the species tree topology (Figure 1e,f). Because discordant pseudoorthologs are difficult to identify—and may introduce both branch length and topological heterogeneity—they are often invoked as the culprits behind counterintuitive results in phylogenetics.

**Figure 1.**
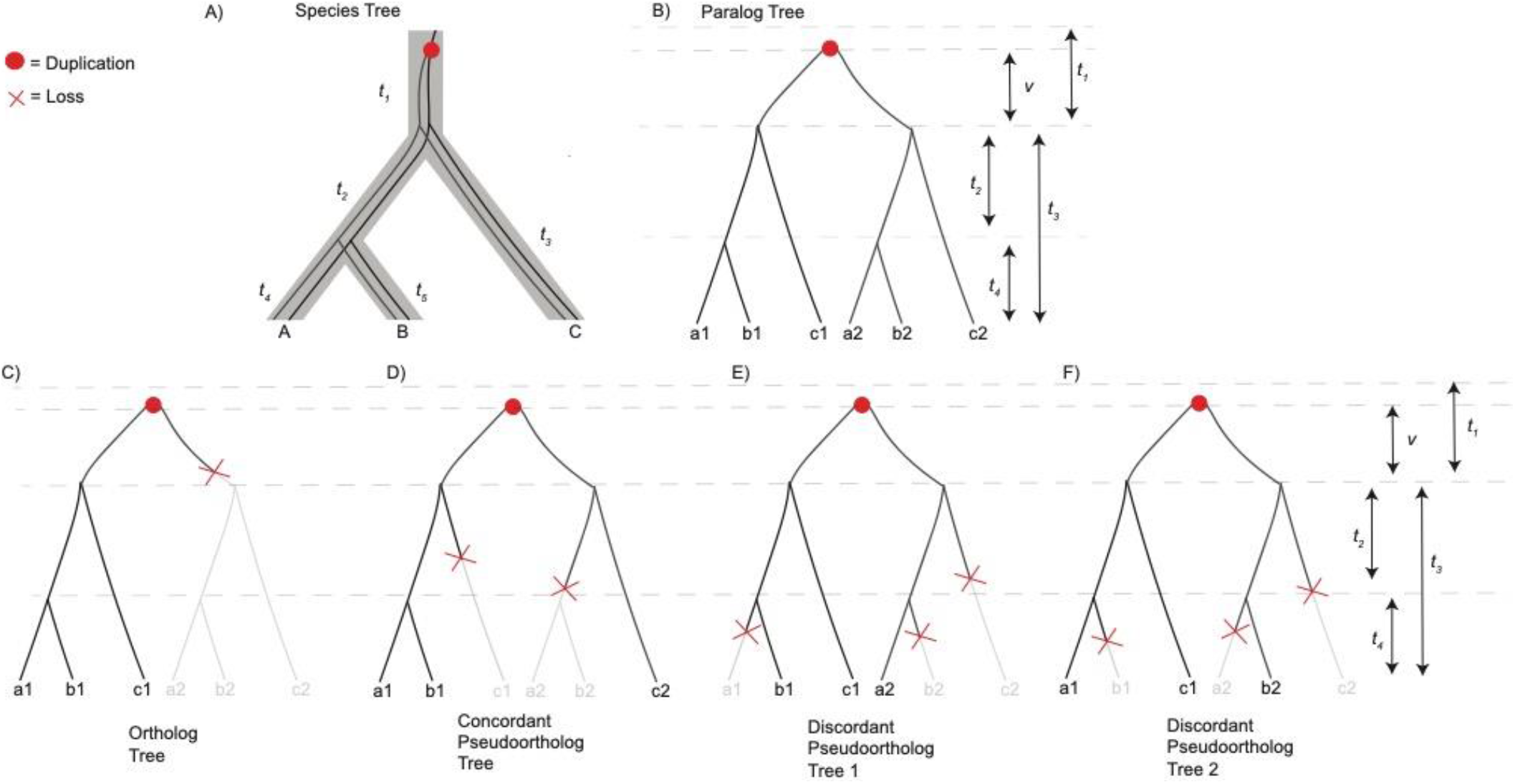
Orthologs and Pseudoorthologs. A) A rooted three-taxon species tree showing the relationships between Species A, B, and C is depicted by the grey outline. Within the tree, the duplication is indicated by a red dot, and the two daughter paralog trees are drawn. B) The full paralog tree depicting relationships between all gene copies. The expected lengths of all branches are shown on the right. C-F) The different relationships possible when a single copy in each species is present. Red Xs indicate loss events. C) Orthologs require at least one loss, D) concordant pseudoorthologs require at least two losses, and E, F) discordant pseudoorthologs require at least three losses.

Multiple studies have attempted to assess the influence of pseudoorthologs on phylogenetic inference, though they have generally done so by comparing results filtered using different ortholog detection methods (3). The results of these analyses have been mixed, with some studies finding substantial differences in inferred species trees (13–15) and others finding minimal differences (14, 16, 17). Furthermore, and in contrast to the long-held opinion that orthologs, not paralogs, should be used to infer species relationships, recent methodological developments explicitly allow for the inclusion of paralogs in phylogenetic inference (reviewed in 18). In particular, quartet-based gene tree methods are robust to the inclusion of paralogs because the concordant topology is expected to be the most common topology, in the limit of a very large number of genes (19–22).

For researchers who wish to only use orthologs for phylogenetic inference, current practices for identifying such genes can be particularly restrictive. This is because excluding paralogs, particularly pseudoorthologs, from phylogenetic datasets is difficult. When extracting putative orthologs, researchers typically rely on graph-based or tree-based approaches (5). Graph-based approaches are the most commonly used, and they rely on the concept of reciprocal best hits (23). These methods assume that the two most closely related homologs between a pair of species should be orthologs. When pseudoorthologs are present they will be reciprocal best hits, despite not having an orthologous relationship, because true orthologs are absent. Tree-based approaches (e.g. 7) are more computationally intensive, but in some cases may be able to identify and exclude pseudoorthologs. In an attempt to completely exclude pseudoorthologs (and paralogs in general) from phylogenetic datasets, researchers often rely on stringent filtering approaches that drastically reduce the amount of data available. For example, Cheon et al. (14) compared two orthology-detection methods on a dataset of mammal transcriptomes: a graph-based approach (6) thought to be susceptible to pseudoorthologs and a tree-based approach (7) thought to be more rigorous. Use of the latter approach reduced their dataset from more than 2000 putative orthologs to only 270. Similarly, Siu-Ting et al. (15) found that stricter filtering to exclude potential pseudoorthologs removed 637 of their 2656 putative orthologs. Thus, a better understanding of when, and how stringently, we should filter our data to remove pseudoorthologs could prevent unnecessary filtering of informative genes from phylogenetic datasets.

Despite long-standing concerns about the effects of pseudoorthologs on phylogenetic inference, no attempt has been made to calculate the probability of pseudoorthologs or to understand the regions of parameter space in which they may be most problematic. Here, we use a stochastic birth-death model to calculate the probabilities and branch lengths of orthologs and pseudoorthologs, including both concordant and discordant pseudoortholog topologies. In what follows, we first describe the model, and then explore regions of parameter space that are most likely to produce pseudoorthologs. We show that the probability of orthologs is high relative to the probability of pseudoorthologs across parameter space, and that the ratio of concordant to discordant topologies is even higher. Our results should reassure researchers using less stringent ortholog filtering methods, greatly expanding the number of genes that can be used for phylogenetics.

## Results

### The Model

To calculate the probabilities of orthologs and pseudoorthologs, we use a stochastic birth-death model (24). Previous work has applied birth-death models to gene trees with the aim of inferring orthology, reconciling gene and species trees, and accurately reconstructing gene trees (25–27). Here, we evaluate a specific case by focusing on a rooted three-taxon species tree, including all scenarios that generate single-copy genes in order to estimate probabilities of orthologs and pseudoorthologs. All calculations assume that there is one gene copy at the beginning of internal branch *t*_*1*_ (Figure 1a). When only a single duplication (and no loss) occurs on this branch, such that two copies exist at the most recent node, we treat each copy independently, generating two “daughter” gene trees (Figure 1b). The independent evolution of each copy means that we can calculate probabilities of further gain and loss on all subsequent lineages, always beginning with a single copy at the base of the daughter gene trees. Since we always begin with a single copy, we use the following equations to calculate the probabilities of transitions along branches, where *λ* is the duplication rate and *μ* is the loss rate. The probability of starting with 1 copy and ending with *n* copies along a branch with length *t* can be calculated as (24):

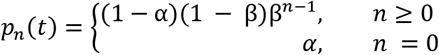

where

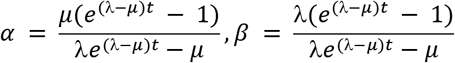

When *λ* = *μ*, we use the following simplification (24):

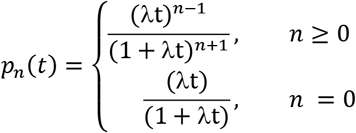

Using the above equations, we can calculate the overall probabilities of different ortholog and pseudoortholog topologies. As an example, consider the concordant pseudoortholog in Figure 1d (see also Supporting Figure S2a). We calculate the overall probability of this topology by multiplying: the probability of transitioning from 1 to 2 copies on branch *t*_*1*_, the probability of transitioning from 1 to 0 copies on branch *t*_*2*_ in one copy and 1 to 0 copies on branch *t*_*3*_ in the other copy, and the probability of no changes on any of the other branches (Appendix A). Note that there are two different paths by which this outcome is achieved, depending on which daughter gene tree losses occur on. Similarly, we can calculate the probability of the type of ortholog shown in Figure 1a by calculating the total probability that there are no transitions on any branch (i.e. that the state is 1 at all nodes; Supporting Figure S1a). In total, we consider six ortholog configurations (Supporting Figure S1) that can each occur in from one to six different arrangements (depending on which exact copies are lost). There are nine concordant pseudoortholog configurations (Supporting Figure S2) and five configurations for each of the two discordant pseudoortholog topologies (Supporting Figure S3), each of which can occur in one to six different arrangements. There are more configurations possible with more ancestral copies, but here we limit the number of copies at the end of branch *t*_*1*_ to 3 (i.e. two duplications on branch *t*_*1*_). While this does not exhaust the possibilities, we verified with simulations in SimPhy (28) that the assumptions made here lead to accurate predictions of the numbers of orthologs, as well as concordant and discordant pseudoorthologs (Supporting Figure S4). Code to calculate the probabilities of orthologs, concordant pseudoorthologs, and discordant pseudoorthologs is shown in Appendix A.

### Probabilities of Orthologs and Pseudoorthologs

We used the model described above to explore how different parameters affected the probabilities of orthologs and pseudoorthologs, including both concordant and discordant topologies. Considering the rates of gene duplication (*λ*) and loss (*μ*), we found that higher rates of each decreased the overall the probability of orthologs (Supporting Figure S5a). The probability of single-copy orthologs decreases because there is a higher chance of duplication and loss events occurring: duplication creates additional copies, while loss means that no copy can be sampled from some species. A similar effect is generated by increasing all branch lengths.

By contrast, the probability of pseudoorthologs is maximized at intermediate values of *λ* and *μ* (Figure 2a), because at least one duplication event and two loss events are required for pseudoorthologs (Figure 1d-f). Values of these parameters that are too high decrease the probability of there being a single copy in each species, though because pseudoorthologs require more losses than gains, slightly higher values of *μ* are possible. Similarly, the probability of pseudoorthologs is maximized at intermediate branch lengths of *t*_*1*_ (Figure 2b) and *t*_*3*_ (Supplementary Figure 5b), because at least one duplication is required on branch *t*_*1*_ and at least one loss is required on branch *t*_*3*_.

**Figure 2.**
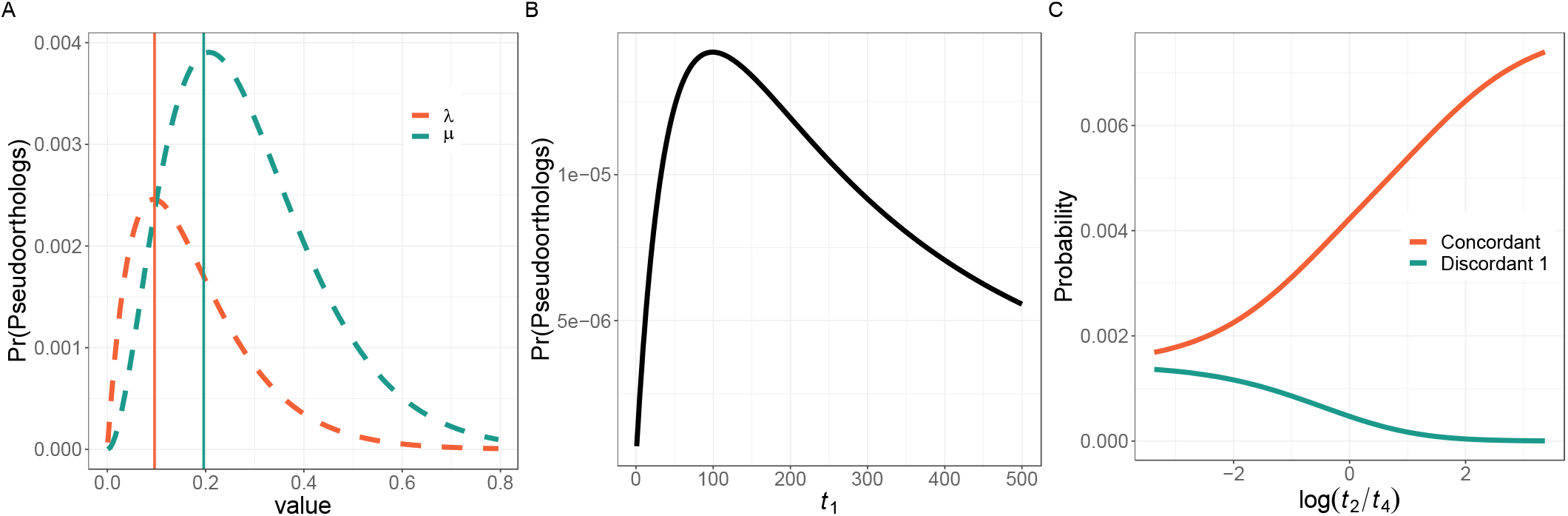
Effects of varying parameters on the probability of pseudoorthologs. A) The unconditional probability of pseudoorthologs is maximized at intermediate values of *μ* and *λ*. B) The unconditional probability of pseudoorthologs is maximized at intermediate values of branch length *t*_*1*_. C) The relative unconditional probabilities of concordant and discordant pseudoorthologs depend on the ratio of branch lengths *t*_*2*_ and *t*_*4*_ (or *t*_*5*_). As *t*_*2*_ gets larger (x-axis becomes more positive) the unconditional probability of concordant pseudoorthologs increases and the unconditional probability of discordant pseudoorthologs decreases. The probability of one discordant pseudoortholog is shown, but the values are identical for the other discordant pseudoortholog.

### Probabilities of Concordant and Discordant Pseudoorthologs

To understand the relative probabilities of concordant and discordant pseudoortholog topologies, we examined many of the same parameters. Increasing *μ* decreases the ratio of concordant pseudoorthologs to discordant pseudoorthologs, because discordant pseudoorthologs require at least three losses while concordant pseudoorthologs can occur with only two losses (Supporting Figure S5c). Increasing *λ* slightly increases the ratio of concordant pseudoorthologs to discordant pseudoorthologs (Supporting Figure S5c), as there are more possible configurations leading to concordant pseudoorthologs than discordant pseudoorthologs when there is more than one duplication event (Supporting Figures S2, S3). Changes to the lengths of branches *t*_*2*_ and *t*_*4*_/*t*_*5*_ affect the relative probabilities of concordant and discordant pseudoorthologs: specifically, as branch *t*_*2*_ gets longer and branches *t*_*4*_/*t*_*5*_ get shorter, concordant pseudoorthologs become more likely and discordant pseudoorthologs become less likely (Supporting Figure 2c). This occurs because concordant pseudoorthologs can be generated either by a loss on *t*_*2*_ or by losses on both *t*_*4*_ and *t*_*5*_ (Figure 1d; Supporting Figures S1a,b), while discordant pseudoorthologs require losses on branches *t*_*4*_ and *t*_*5*_ (Figure 1e-f; Supporting Figure S3). Both scenarios additionally require losses on *t*_*3*_, and so the length of *t*_*3*_ does not affect their relative frequencies. Note that results from the model presented here are also supported by results from simulations (Supporting Figure S4).

We further explored the probabilities of all events conditional on a single copy being present in each species. These calculations directly address the chance that orthologs are mistaken for pseudoorthologs: the conditional probabilities represent the fraction of all single-copy genes that are orthologs, concordant pseudoorthologs, or discordant pseudoorthologs. We explored two general regions of parameter space, representing the range of values of *λ* and *μ* observed in empirical datasets: 0.002 and 0.005 per million years (29). We considered a long length of branch *t*_*3*_ (198.9 million years) across a range of lengths for branches *t*_*1*_, *t*_*2*_, *t*_*4*_, and *t*_*5*_. The large value of *t*_*3*_ mirrors a potentially difficult region of tree space (see next section), coupled with moderate and high rates of duplication and loss.

The conditional probability of orthologs given that a single copy is present in each species is very high when rates of duplication and loss are moderate (0.002, Figure 3a; minimum conditional probability of orthologs = 0.955), and is moderately high even when rates of duplication and loss approach the highest observed in empirical datasets (Figure 3c; minimum conditional probability = 0.711). Furthermore, the ratio of concordant to discordant topologies is very high when duplication and loss rates are moderate (Figure 3b; minimum = 76.7:1) and is still rather high even when rates of duplication and loss are high (Figure 3d; minimum = 8.4:1). These ratios include both orthologs and concordant pseudoorthologs in the “concordant” category. Note again that we chose *t*_*3*_ to mirror the most problematic regions of parameter space for these results; Supporting Figure S6 shows results for different values of *t*_*3*_, confirming the impression that the scenario shown here in the main text is a worst-case scenario with regards to this branch length.

**Figure 3.**
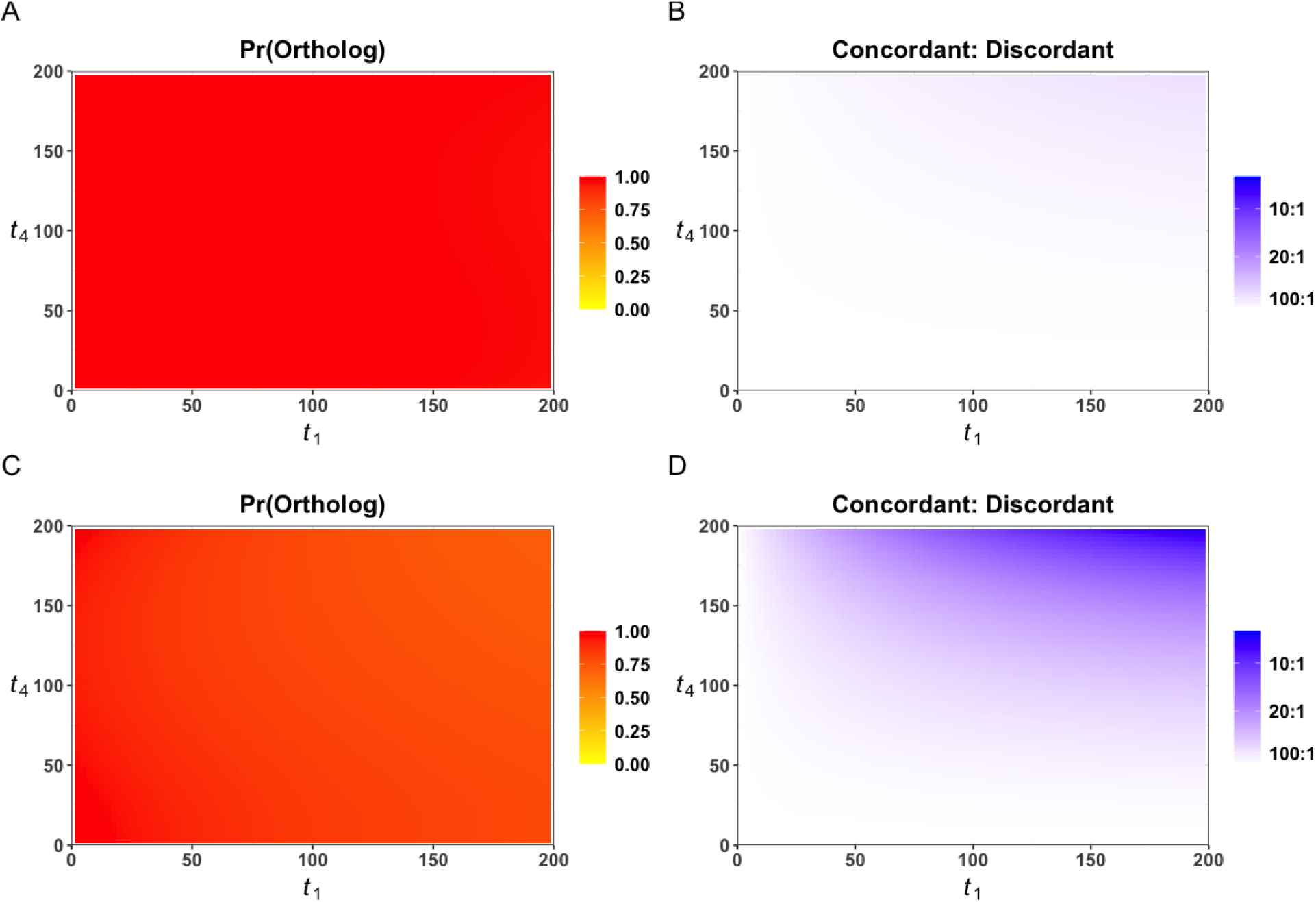
Probabilities of orthologs, pseudoorthologs, and discordance. Here, branch length *t*_*3*_ = 198.9 mya. Branch length *t*_*1*_ varies from 0.0001 to 200 mya, while branch length *t*_*4*_ varies from 0.0001 to 198.8 mya; branch length *t*_*2*_ is constrained such that the sum of *t*_*2*_ and *t*_*4*_ equals *t*_*3*_. A) The probability of orthologs given that a single copy is present in each species, with moderate rates of duplication and loss (*λ*=0.002 per my, *μ*=0.002 per my). B) The ratio of the concordant topology (orthologs and concordant pseudoorthologs) to one discordant topology given that a single copy is present in each species, with moderate rates of duplication and loss (*λ*=0.002, *μ*=0.002). C) The probability of orthologs given that a single copy is present in each species with high rates of duplication and loss (*λ*=0.005, *μ*=0.005). D) The ratio of the concordant topology (orthologs and concordant pseudoorthologs) to one discordant topology given that a single copy is present in each species with high rates of duplication and loss (*λ*=0.005, *μ*=0.005).

Notably, the probability of either of the two discordant pseudoorthologs can never exceed the probability of the concordant pseudoortholog, because it is always possible to generate a concordant pseudoortholog with the same number of duplication and loss events (and often fewer events). For example, a discordant pseudoortholog can be generated by a duplication on branch *t*_*1*_, a loss on branches *t*_*4*_ and *t*_*3*_ in one copy, and a loss on branch *t*_*5*_ in the other copy (Figure 1e). A concordant pseudoortholog could also be generated by this pattern, as long as the losses on branches *t*_*4*_ and *t*_*5*_ occurred in the same copy, while the loss on branch *t*_*3*_ occured in the other copy (Supporting Figure S2b). In reasonable regions of parameter space, the probability of concordant pseudoorthologs is much higher than the probability of either discordant pseudoortholog because most concordant pseudoorthologs require one fewer loss event (i.e. the scenario shown in Figure 1d; Supporting Figure S2, S3). Moreover, the probabilities of the two discordant topologies are always equal, as these rely on the same events on the same branches, and only differ in terms of which branches are lost from which copy. Thus, one never expects either discordant topology to be significantly more frequent than the other. Finally, note again that in Figure 3 we are showing the probabilities of orthologs and pseudoorthologs conditional on sampling a single copy per species. However, the absolute probability of sampling a single copy per species at all is lowest in the regions of parameter space that maximize the probability of pseudoorthologs and discordant topologies (Supporting Figure S7).

### Worst-case scenarios

In order to find the species tree topologies most prone to producing pseudoorthologs (especially discordant ones), we searched parameter space for such trees. Using a straightforward optimization procedure (Methods), we found the regions of parameter space that maximize the conditional probabilities of pseudoorthologs, while minimizing the ratio of concordant to discordant topologies (Figure 4; Tables S1, S2). The maximum conditional probability of pseudoorthologs that we found was 0.285, and this value was only obtained with values of *λ* and *μ* higher than observed in most empirical systems (29). The highest conditional probability of either of the two discordant pseudoorthologs observed was 0.095, and again this involved high values of *λ* and *μ* (Figure 4).

**Figure 4.**
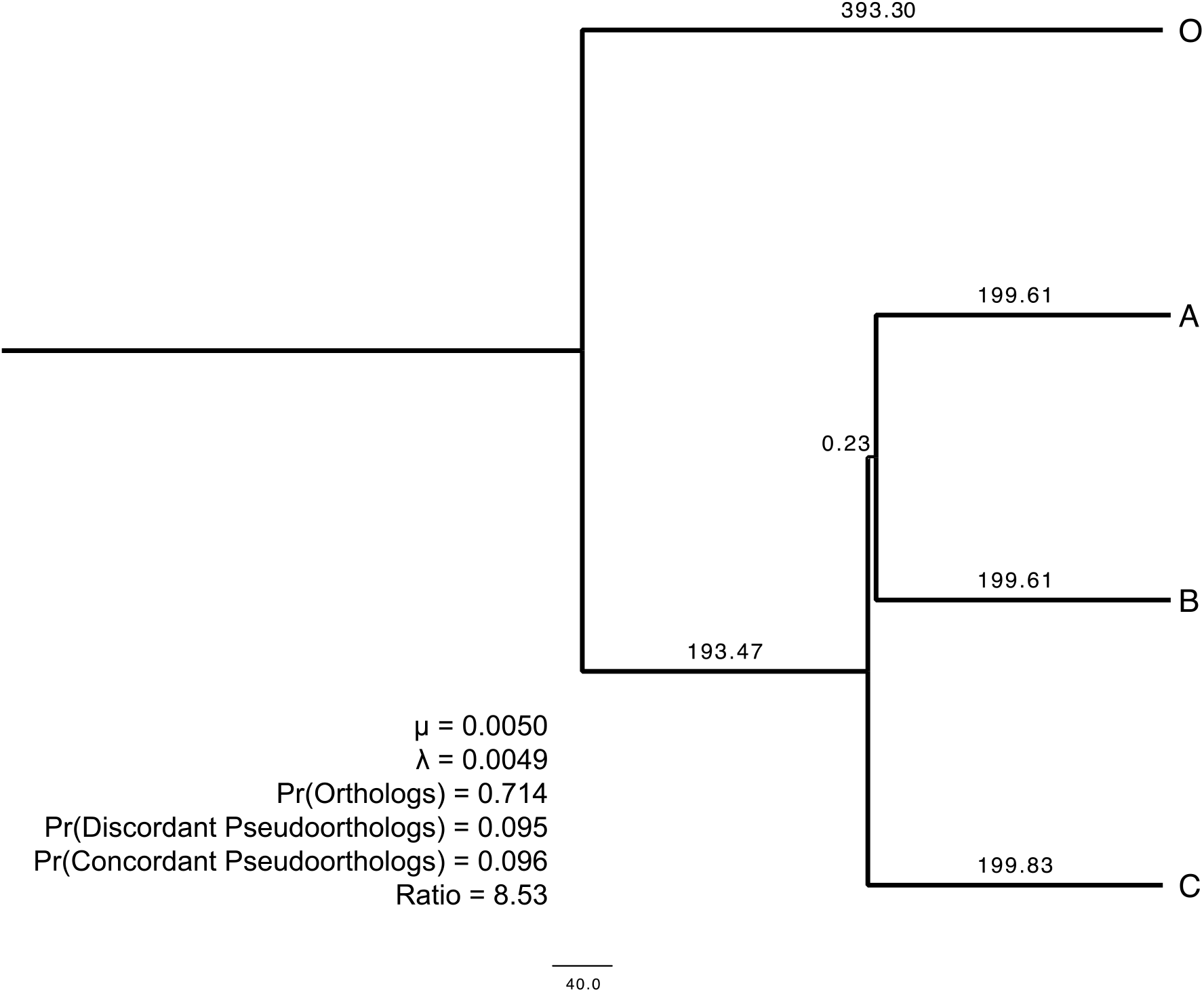
The species tree and parameters that minimize the ratio of the concordant topology to the two discordant topologies. Discordant probabilities and ratios refer only to discordant pseudoortholog 1, but the probabilities of the two discordant pseudoorthologs are equal. Probabilities are conditional on sampling a single copy per species. To facilitate visualization, the internal branch uniting Species A and B is not drawn to scale.

The minimum ratio of the concordant topology to either of the two discordant topologies was 8.5. This suggests that, even in the most problematic regions of parameter space, discordant pseudoorthologs will comprise fewer than 10% of all single-copy genes.

Equally importantly, our results show that the ratio of concordant to discordant topologies is lowest in a region of tree space in which discordance due to ILS is also likely to be a concern (Figure 4)— when the internal branch of a three-species tree is very short. If we take units in millions of years and assume a species with a generation time of 29 years and an effective population size of 10,000, then the probability of either discordant topology for the worst-case species tree under the multispecies coalescent model is 0.225. By comparison, the conditional probability of either discordant topology under our model of duplication and loss in this same area of parameter space is 0.095, a value more than two times lower. Additionally, this region of parameter space involves very long branch lengths for *t*_*1*_, *t*_*3*_, and *t*_*4*_/*t*_*5*_ (nearly 200 million years) and high rates of duplication and loss. In such species trees pseudoorthologs are therefore not likely to be the biggest impediment to phylogenetic inference.

We also explored regions of parameter space that maximized the absolute probabilities of pseudoorthologs and discordant pseudoorthologs, rather than the probabilities conditional on a single copy per species (Supporting Table S2). The most notable difference was in the branch lengths that maximized the probability of pseudoorthologs. When absolute probabilities are considered, a long internal branch *t*_*2*_ and short terminal branches *t*_*4*_ and *t*_*5*_ maximize this probability because they maximize the probability of the concordant pseudoortholog (Supporting Table S2). However, when conditional probabilities are considered, a shorter internal branch *t*_*2*_ maximizes the probability of pseudoorthologs because, coupled with longer branches *t*_*4*_ and *t*_*5*_, a shorter branch *t*_*2*_ decreases the probability of orthologs.

### Extending the model to larger trees

Thus far we have considered the probability of orthologs and pseudoorthologs in a three-taxon species tree. While we might intuitively expect that the addition of more species would lower the relative probability of pseudoorthologs (because more losses would be required to mimic orthologs), we carried out additional analyses to evaluate slightly larger trees by adding a single extra taxon sister to Species A or Species C. These results support our prediction: adding leaves decreases the probability of all types of single-copy genes, including orthologs and concordant pseudoorthologs (Supporting Figure S8a, S8b). However, adding leaves disproportionately decreases the probability of discordant pseudoorthologs, particularly when branches are added as sister to Species A (Supporting Figure S8c, S8d). This outcome occurs because discordant pseudoorthologs require losses on branches *t*_*4*_ and *t*_*5*_, and, if these losses do not occur before the split between the two sister branches including Species A, then the number of losses required increases by one. While the same is true when a branch is added sister to Species C, since branch *t*_*3*_ is longer than branches *t*_*4*_ and *t*_*5*_, there is more time for the loss to occur prior to the added speciation event. Overall, these limited extra analyses indicate that results for a three-taxon tree represent a worst-case scenario for confusing orthologs with pseudoorthologs.

### Expectations for branch lengths of pseudoorthologs

In addition to differing topologically from the species tree, pseudoorthologs differ from the species tree in terms of branch lengths. For orthologs, the internal branch is length *t*_*2*_; this branch determines the phylogenetic signal within each gene tree, and is the focus here. For the simplest concordant pseudoortholog (Figure 1d), the internal branch length is equal to the sum of *t*_*2*_ and the time until the duplication event occurs in branch *t*_*1*_ (*v* in Figure 1), while for the simplest discordant pseudoortholog the internal branch length is equal to *v* (Figure 1e,f). Furthermore, internal branch lengths for the two discordant pseudoorthologs are always equal.

The value of *v* is the expected time back to the duplication event on *t*_*1*_ conditional on a duplication event occurring on branch *t*_*1*_. Because waiting times for events in the birth-death process are also exponentially distributed (30), we can use a model similar to that for the multispecies coalescent (31) to calculate times here. To find the expectation for *v*, we need only convert from the coalescent units used in ref. (31), to duplication units, where one coalescent unit is equal to 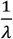. These considerations lead to the following expectation:

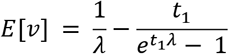

Although we have conditioned on a duplication event occurring in branch *t*_*1*_, we have not conditioned on other events. For example, we have not conditioned on the absence of any subsequent duplication events or a subsequent loss on branch *t*_*1*_. Despite this, the expected branch lengths are a close match for simulated branch lengths (Supporting Figure S9), and thus should provide insights into the internal branch lengths of pseudoorthologs.

Together, the results here demonstrate that the expected internal branch length for concordant pseudoorthologs is always longer than the expected branch length for discordant pseudoorthologs, by the length of the internal branch *t*_*2*_. Thus, even when pseudoorthologs are present, the total expected branch length supporting the concordant topology should exceed the expected branch length supporting the discordant topology. In other words, there is more phylogenetic signal in concordant trees than discordant ones.

## Discussion

Pseudoorthologs play an interesting role in phylogenetics: feared for their possible detrimental effects on species tree inference, a number of methods have been created solely to ensure that they are completely excluded from the data used as input to tree-building software. Our results suggest that the use of stringent methods for filtering pseudoorthologs is unnecessary, as they are unlikely to mislead phylogenetic inferences. We find that pseudoorthologs are rare overall (Figure 3a), and that pseudoorthologs with discordant topologies are expected to be much less common than genes with concordant topologies (Figure 3b). Regardless of the particular method used to identify single-copy orthologs, discordant pseudoorthologs are unlikely to be mistakenly sampled; thus, they are unlikely to pose a challenge to phylogenetics.

Even in the most problematic regions of parameter space considered here, pseudoorthologs are unlikely to mislead topological inferences. First, orthologs are still substantially more likely than pseudoorthologs, and the concordant topology is still more than 8X as likely as either of the two discordant topologies. Second, topological heterogeneity is recognized as common across the tree of life due to many processes (32), and in this particular region of parameter space we expect discordance to be high due to incomplete lineage sorting (Figure 4). Since discordance is likely to be high irrespective of the presence of pseudoorthologs, the use of methods that are robust to ILS (e.g. ASTRAL 33, 34) should be of utmost importance. Moreover, our modelling results corroborate previous findings that quartet methods such as ASTRAL are statistically consistent under a model of gene duplication and loss (20, 21). Such methods rely on the fact that, for a rooted three-taxon species tree, the concordant topology is always the most frequent, which is exactly what we find here among all single-copy genes. Additionally, the probabilities of the two discordant topologies are always equal, suggesting that methods for species tree inference (35) and introgression tests (36, 37) based on symmetry in number of the two discordant topologies should not be misled by the inclusion of pseudoorthologs. Thus, even in the most problematic regions of parameter space, quartet-based methods should not be misled by the presence of pseudoorthologs. Finally, in these regions of parameter space—and when there are more than three species in a tree—single-copy genes shared across all species are particularly rare (e.g. Supporting Figure S7b). Thus, we expect that researchers would be unable to sample many single-copy genes in such cases, and should therefore consider explicitly including paralogs in their dataset to gain more phylogenetic markers (18).

Our model and simulations have not considered every scenario, nor every effect of pseudoorthologs. Polyploidy is a special case of gene duplication and loss in which the entire genome is duplicated (38), but we believe that our results should still apply. When the considered taxa are autopolyploids, or polyploid taxa for which both sub-genomes came from the same species, then we need only condition on the polyploidy event having occurred at some point on branch *t*_*1*_ for our model to apply. While the guaranteed duplication on this branch decreases the overall probability of orthologs and increases the overall probability of pseudoorthologs (though many paralogs will be lost before the first speciation event (39)), there is no effect on the relative probability of discordant and concordant topologies. Thus, even in autopolyploids, the probability of either discordant pseudoortholog can never exceed the probability of the concordant pseudoortholog. Allopolyploids are polyploid taxa in which each sub-genome comes from a different species (38). Although there can be cases where paralogs from the different parental species are lost in a biased fashion (e.g. 40), this should not affect the probability of producing discordant pseudoorthologs since these still require losses from each set of copies (Figure 1e,f). The main consequence of biased loss is that one set of orthologous species relationships will be retained over the other (cf. 41). Of course, even more so in polyploids than in other taxa, excessive filtering to remove putative paralogs will decrease the amount of available data, and researchers would likely benefit from using paralogs for phylogenetic inference.

In addition to changes in gene tree topologies, pseudoorthologs have different branch lengths than orthologs. Concordant pseudoorthologs are expected to have the longest internal branch lengths, with the expected branch length converging on *t*_*2*_ + 1/*λ* (the latter term representing the expected time to the duplication event) as the length of branch *t*_*1*_ increases (Figure 1d; (31)). Discordant pseudoorthologs will have longer terminal branch lengths (Figure 1e, f), but a shorter internal branch length. The expected internal branch length of discordant pseudoortholog will converge to 1/*λ* as the length of branch length *t*_*1*_ increases, which may be either shorter or longer than the internal branch length *t*_*2*_ of true orthologs. Since concordant pseudoorthologs are never expected to occur at a lower frequency than either of the discordant pseudoorthologs—and the internal branch is always longer by *t*_*2*_—the total expected internal branch length supporting the true topology should always exceed that supporting either discordant topology. With enough data, these results suggest that concatenation-based methods are also unlikely to be misled by pseudoorthologs. Although pseudoorthologs could lead to biased branch length estimates, their rarity across most of parameter space (Figure 3a) should minimize their effects on estimates of branch lengths.

Based on the results presented here, pseudoorthologs are unlikely to be the frequent cause of problems in phylogenetic inference. How then can we explain previous results that demonstrate the importance of filtering datasets for orthologs? Some studies have found differences in trees inferred from datasets filtered using different ortholog detection methods (e.g. 14, 15). Of course, more stringent filtering may remove problematic sequences other than paralogs, for example alternative isoforms or error-prone sequences. Also notably, in these studies paralogs included in datasets of putative single-copy orthologs may not be true pseudoorthologs. In large genomic and transcriptomic datasets, putative pseudoorthologs may rather be paralogs for which, for technical reasons, different copies were assembled in each species. However, even in the extreme case, where a single paralog is sampled at random from each species, quartet-based methods appear to perform well when enough data is available (19, 20). This result is also supported by the work presented here: in our model, *μ* need not represent only the rate of true loss. Rather, we can consider *μ* to be the rate of both true loss and apparent loss due to sampling error. Although in this case *μ* may be much higher, the finding that the probability of a discordant topology cannot exceed that of the concordant topology still holds.

Overall, our results suggest that the stringent filtering often adopted to keep datasets free from pseudoorthologs is unnecessary and removes valuable information from phylogenetic datasets. By embracing the rarity of pseudoorthologs, and the fact that, even when present, they are unlikely to mislead phylogenetic inference, researchers could increase the number of genes used often by an order of magnitude (e.g. 14). With small numbers of genes, even minor issues with dataset compilation (including the inclusion of pseudoorthologs) may play a disproportionate role in phylogenetic inference (e.g. 42). However, as the total number of genes used increases, the likelihood of pseudoorthologs misleading phylogenetic inference decreases, as speciation events should be represented far more often in these datasets than duplication events. Thus, by relaxing filtering thresholds and using more genes for inference, researchers reduce the chances of their results being misled by a small number of problematic genes.

## Materials and Methods

### Finding scenarios that maximize discordance

To better understand the best and worst-case scenarios with regards to pseudoorthologs, we searched for regions of parameter space that a) maximized the probability of pseudoorthologs conditional on a single copy per species, b) maximized the probability of discordant pseudoorthologs conditional on a single copy per species, and c) minimized the ratio of concordant topologies to discordant topologies (Supporting Table S1). We also explored scenarios that maximized the absolute probabilities of pseudoorthologs and discordant pseudoorthologs (Supporting Table S2). We set bounds on all parameters (Supporting Table S3), and constrained the species tree to be ultrametric by setting *t*_*5*_ equal to *t*_*4*_ and requiring that *t*_*4*_+ *t*_*2*_ = *t*_*3*_. We changed each parameter in turn, increasing or decreasing the value at random by a value chosen from a uniform prior distribution U(0.000001, 0.001) for *μ* and *λ*, and U(0.0001, 20) for branch lengths. For each optimization, we accepted each change if it increased the probability (or decreased the ratio), and accepted the change one percent of the time if it decreased the probability (or increased the ratio). For each parameter we performed 100 optimization steps. We visited the parameters in the order: *μ*, *λ*, *t*_*1*_, *t*_*3*_, and *t*_*2*_. We repeated this procedure ten times.

### Simulations in SimPhy

To evaluate whether our assumptions led to accurate predictions, we compared the exact probabilities calculated according to the equations above to the proportions of each scenario observed in simulations in SimPhy (28). We calculated the proportion of observed orthologs, concordant pseudoorthologs, and discordant pseudoorthologs by evaluating topology and branch lengths. If our assumptions are reasonable, we expect a 1:1 relationship between calculations and observations. We drew 100 sets of parameters from uniform priors (Supporting Table S4), and performed 10,000 simulations under each set of parameters in SimPhy to calculate the proportions of observed orthologs, concordant pseudoorthologs, and discordant pseudoorthologs.

We also used simulations to evaluate whether our assumptions about branch lengths led to accurate predictions. We focused on the scenarios from Figure 1d and Figure 1e. We calculated the observed and expected lengths of the internal branch lengths. We drew 50 sets of parameters from uniform priors. All priors were the same as in Supporting Table S4 except *μ* and *λ* were drawn from U(0.004, 0.005) priors to ensure more pseudoorthologs. We performed 10,000 simulations under each set of parameters, and calculated the average internal branch lengths from those simulations that produced trees matching the concordant pseudoortholog in Figure 1d or the discordant pseudoortholog in Figure 1e.

### Expanding the model to larger trees

The above calculations pertain to rooted trees with three leaves (A, B, and C). In principle, these calculations should extend to larger trees, although the number of relevant scenarios quickly becomes prohibitively large when trying to calculate probabilities on very large trees. We extended our calculations to trees with one extra leaf on A, B, or C to evaluate the effects of adding leaves to the tree. We then calculated the probabilities of orthologs, concordant pseudoorthologs, discordant pseudoorthologs, and the ratio of concordant to discordant topologies while varying the length of the additional branches.

## Acknowledgments

We thank Rafael Guerrero for helpful discussion. This work was supported by a National Science Foundation postdoctoral fellowship to MLS (DBI-2009989) and an NSF grant to MWH (DEB-1936187).

