## Supporting Information for "The frequency and topology of pseudoorthologs"

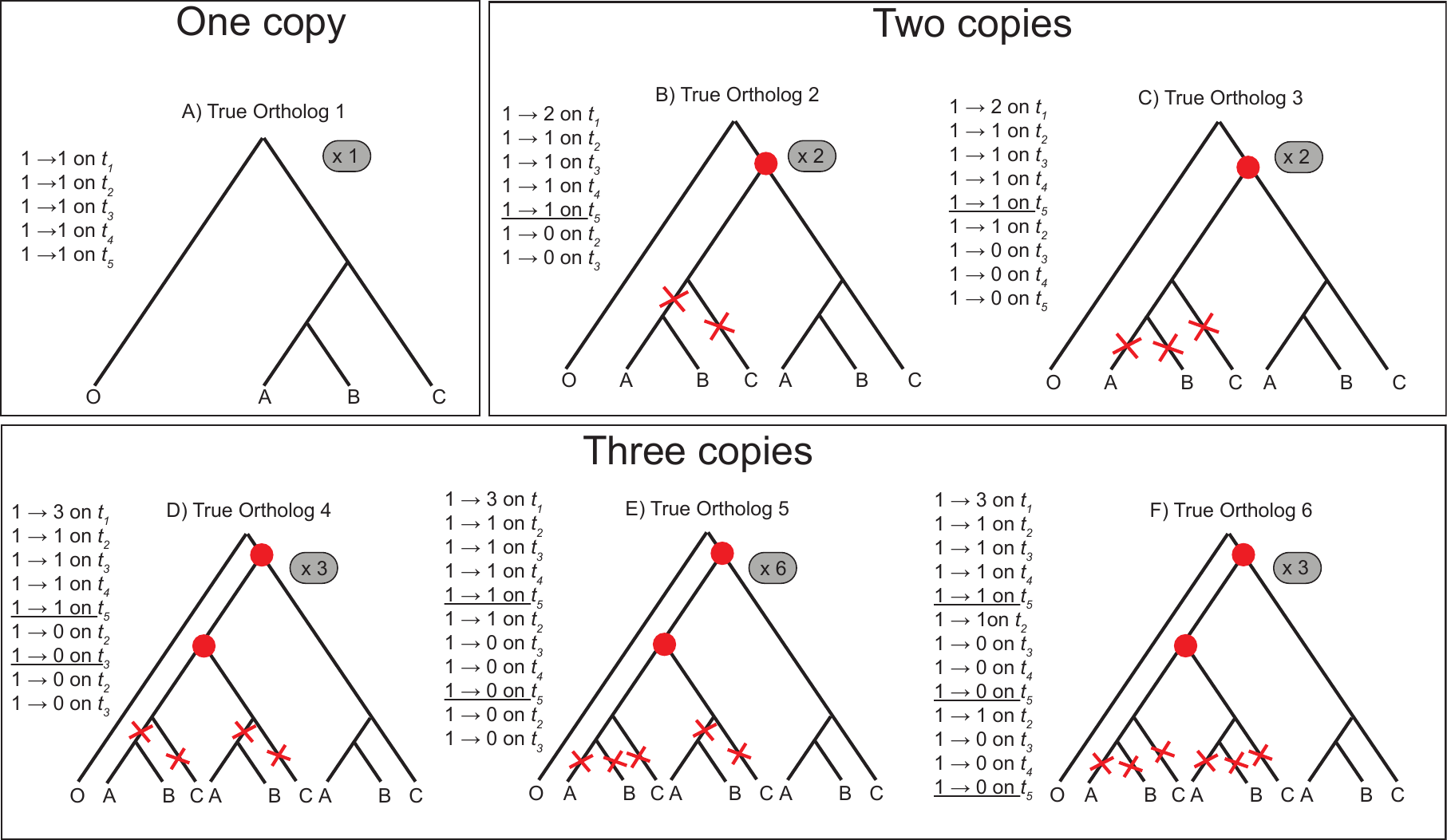
**Fig. S1.** Orthologs considered. Red dots indicate duplication events, and red Xs indicate loss events. The transitions necessary to observe each configuration are listed on the side of each panel. The number of copies at the bottom of branch *t_1_* are one (panel A), two (panels B, C), or three (panels D, E, F). The ‘x 2’ or ‘x 6’ indicates the number of potential arrangements considered for each scenario. For example, for true ortholog 2, the losses could occur in either one of the two copies.

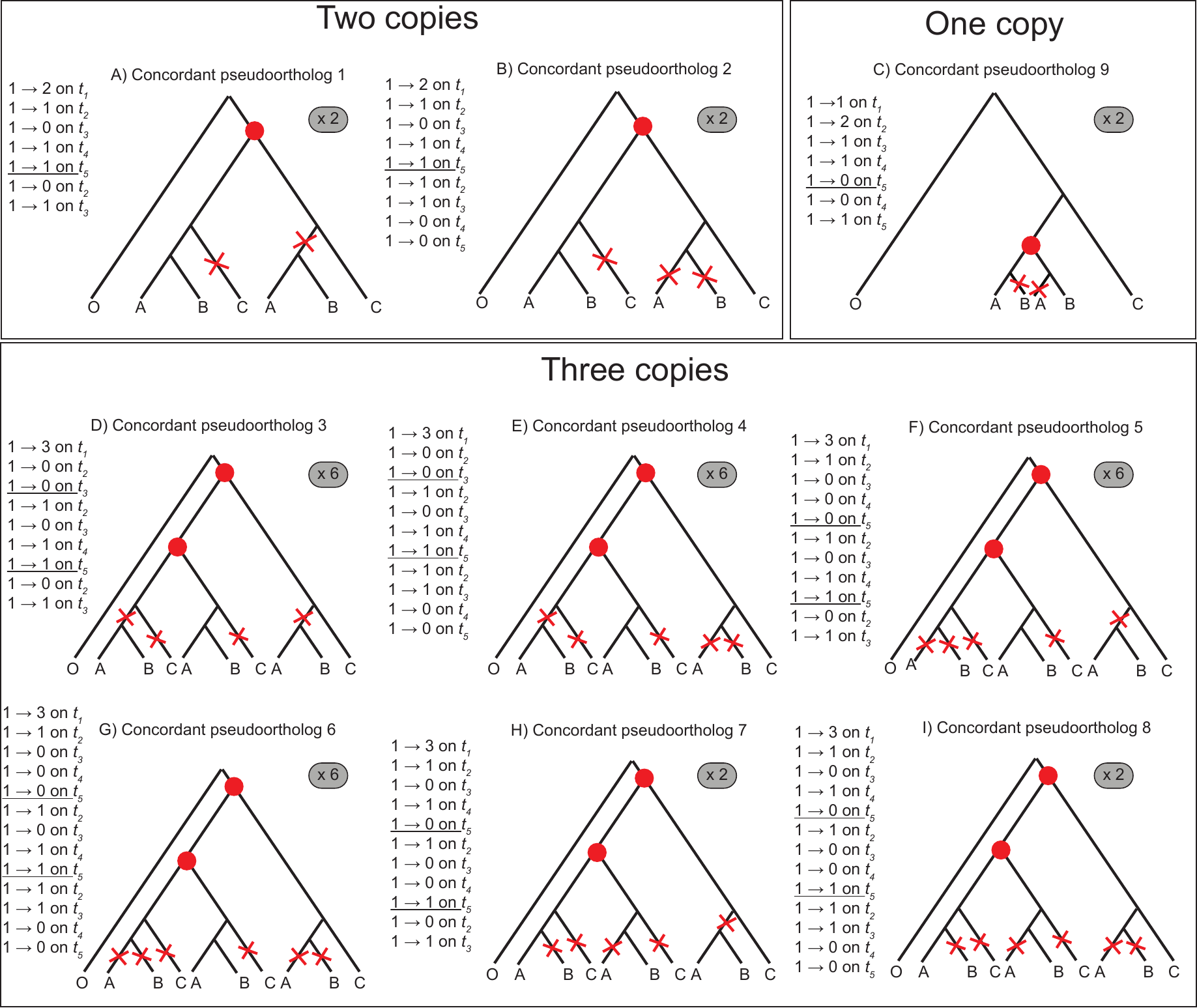
Fig. S2. Concordant pseudoortholog scenarios considered. Red dots indicate duplication events, and red Xs indicate loss events. The transitions necessary to observe each configuration are listed on the side of each panel. The number of copies at the bottom of branch *t_1_* are one (panel C), two (panels A, B), or three (panels D-I). The ‘x 2’ or ‘x 6’ indicates the number of potential arrangements considered for each scenario.

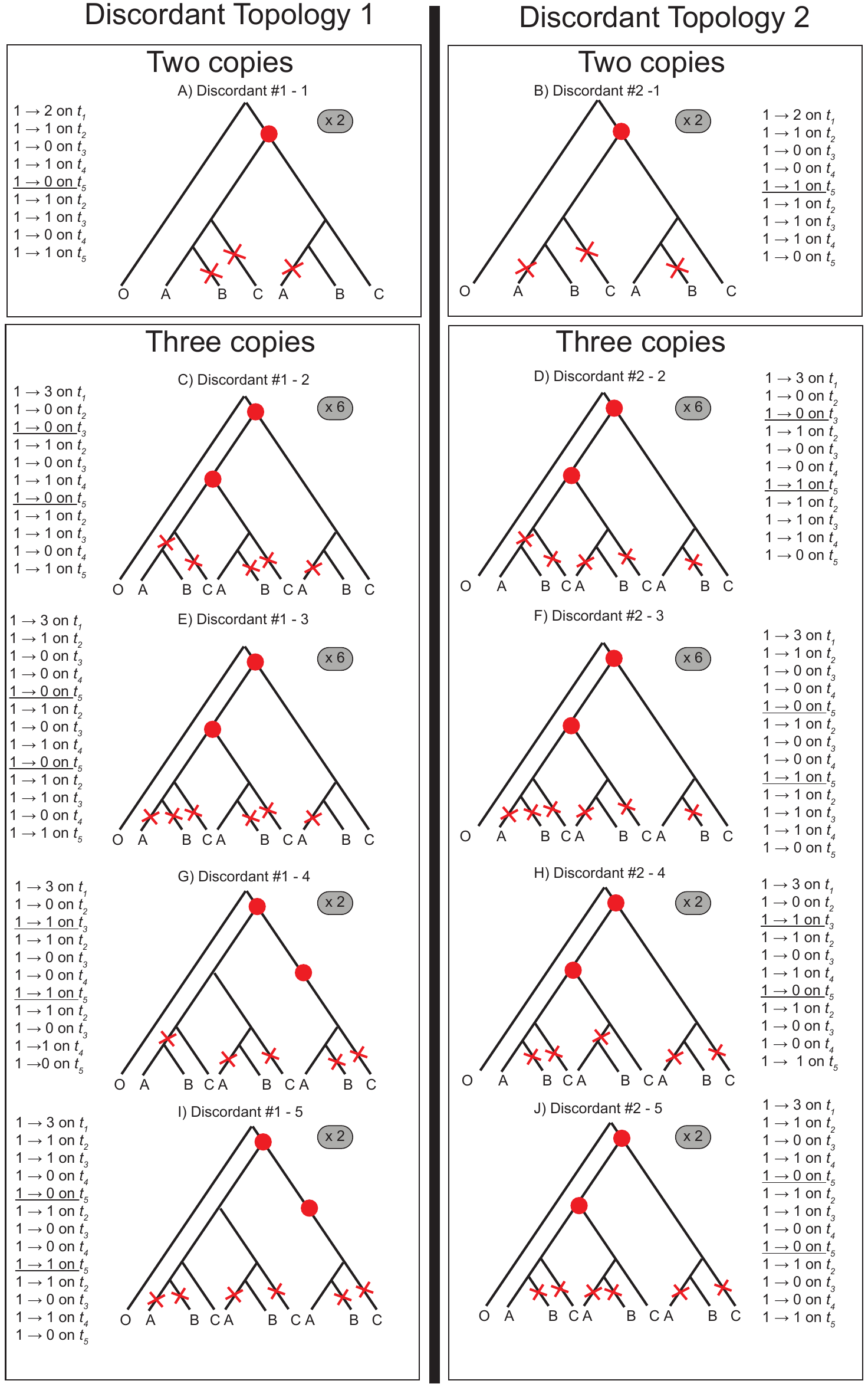

Fig S3. Discordant pseudoortholog scenarios considered. Red dots indicate duplication events, and red Xs indicate loss events. The transitions necessary to observe each configuration are listed on the side of each panel. The number of copies at the bottom of branch *t_1_* are two (panels A, B) or three (panels C-J). The ‘x 2’ or ‘x 6’ indicates the number of potential arrangements considered for each scenario, with the two discordant topologies considered separately in each column.

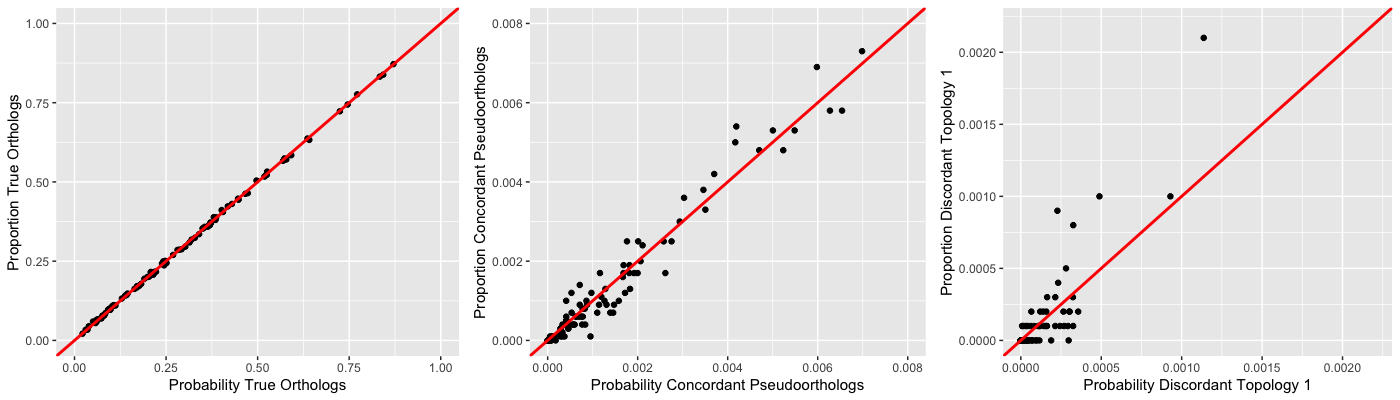
­­­Fig S4. Results of simulations. Red lines indicate the predicted 1:1 relationship between expected probabilities and observed probabilities from simulations. A) The proportion of true orthologs in simulations versus the probability of orthologs under the model. B) The proportion of concordant pseudoorthologs versus the probability of concordant pseudoorthologs under the model. C) The proportion of discordant pseudoortholog 1 versus the probability of discordant pseudoortholog 1 under the model. Note that there are few events in this last category, as the probability of discordant pseudoorthologs is always very low.

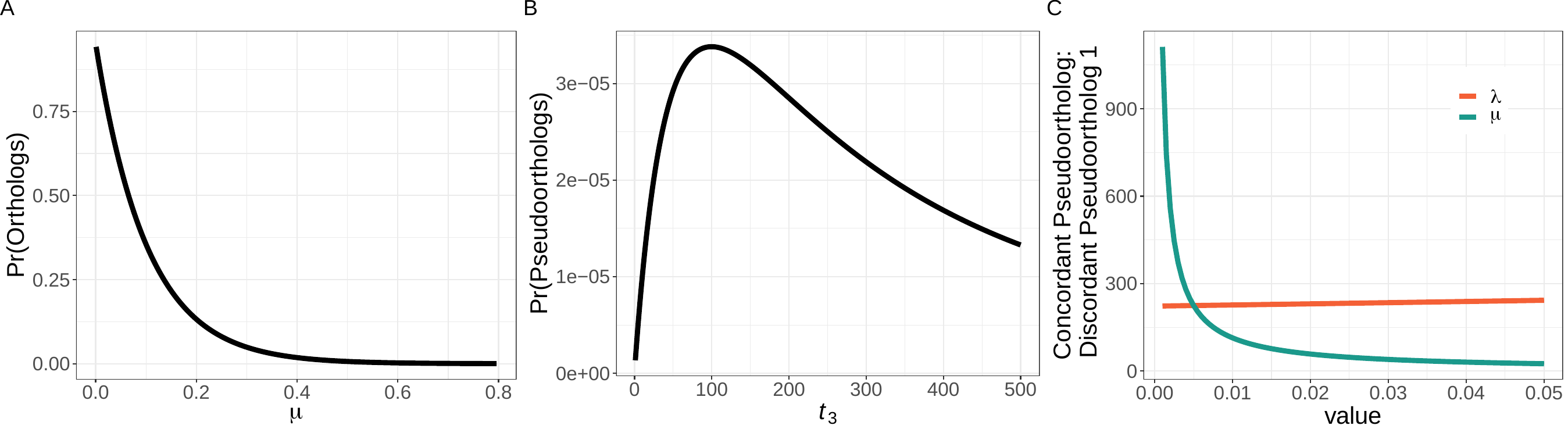

Figure S5. A) The absolute probability of orthologs at values of μ ranging from 0.001 to 0.8.(*t_1_*=5*_,_ t_2_*=1*_,_ t_3_*=2*_,_ t_4_*=1*_,_ t_5_*=1, λ=0.005). Here, we show only the results of varying μ because the results of varying λ are identical. B) The absolute probability of pseudoorthologs at value of *t_3_* ranging from 1 to 500 (λ=0.005, μ=0.005, *t_1_*=5*_,_ t_2_*=1*_,_ t_4_*=1*_,_ t_5_*=1. C) The ratio of the concordant pseudoortholog to one discordant pseudoortholog at values of λ and μ ranging from 0.001 to 0.05 (*t_1_*=5*_,_ t_2_*=1*_,_ t_3_*=2*_,_ t_4_*=1*_,_ t_5_*=1). The green line shows the ratio of the concordant pseudoortholog to one discordant pseudoortholog when μ varies and λ is held constant at 0.005, while the orange line shows the ratio of the concordant pseudoortholog to one discordant pseudoortholog when λ varies and μ is held constant at 0.005.

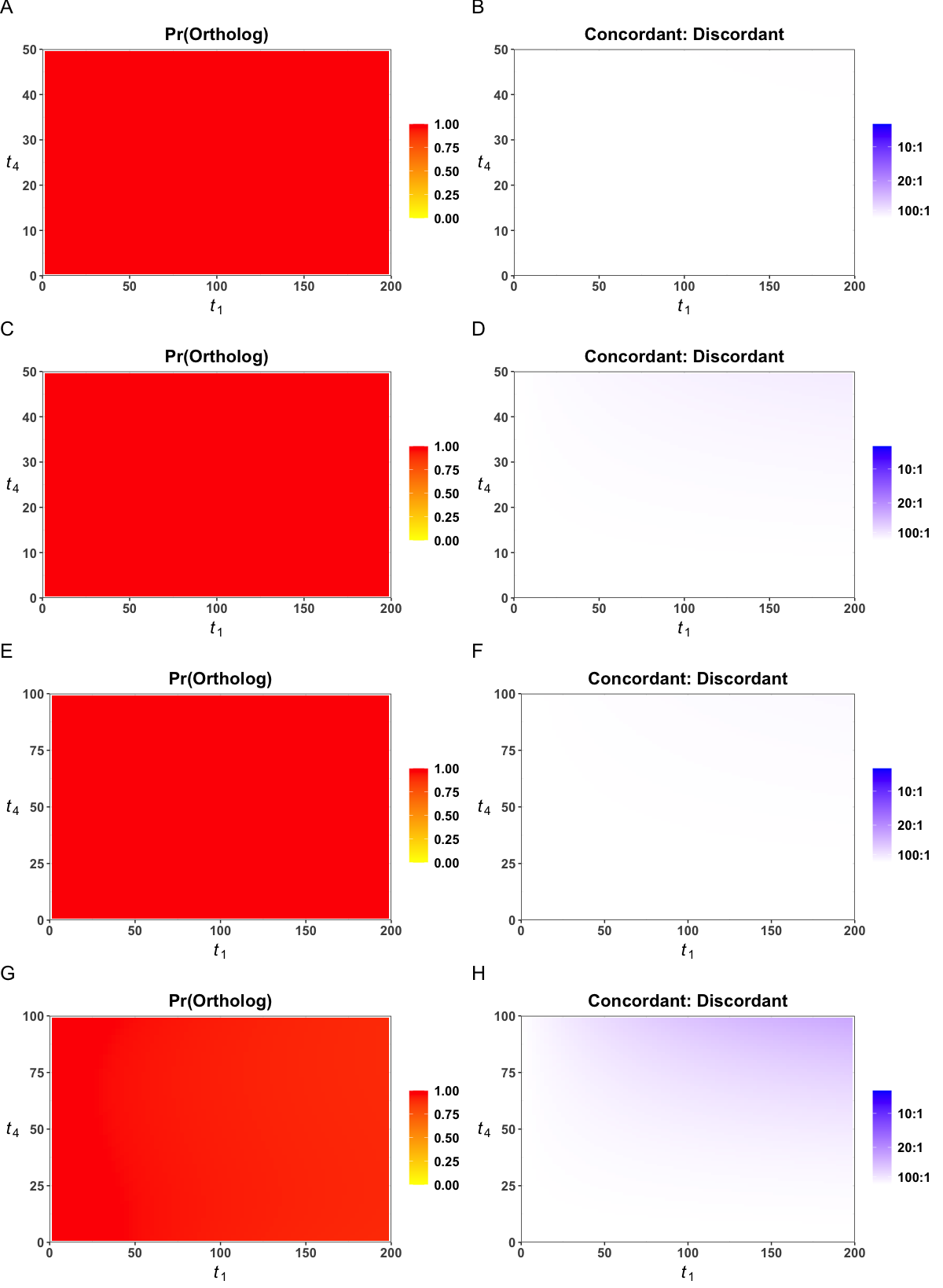

**Fig S6.** Probabilities of orthologs, pseudoorthologs, and discordance conditional on a single copy per species. Branch length *t_1_* varies from 0.0001 to 200 mya, while branch length *t_4_* varies from 0.0001 to *t_3_* -0.1; branch length *t_2_* is constrained such that the sum of *t_2_* and *t_4_* equals *t_3_*. A) The conditional probability of orthologs (λ=0.002 per my, μ=0.002 per my, *t_3_* =50). B) The ratio of the concordant topology to one discordant topology (λ=0.002, μ=0.002, *t_3_* =50). C) The conditional probability of orthologs (λ=0.005, μ=0.005, *t_3_* =50). D) The ratio of the concordant topology to one discordant topology (λ=0.005, μ=0.005, *t_3_* =50). E) The conditional probability of orthologs (λ=0.002 0, μ=0.002, *t_3_* =100). F) The ratio of the concordant topology to one discordant topology (λ=0.002, μ=0.002, *t_3_* =100). C) The conditional probability of orthologs (λ=0.005, μ=0.005, *t_3_* =100). D) The ratio of the concordant topology to one discordant topology (λ=0.005, μ=0.005, *t_3_* =100).

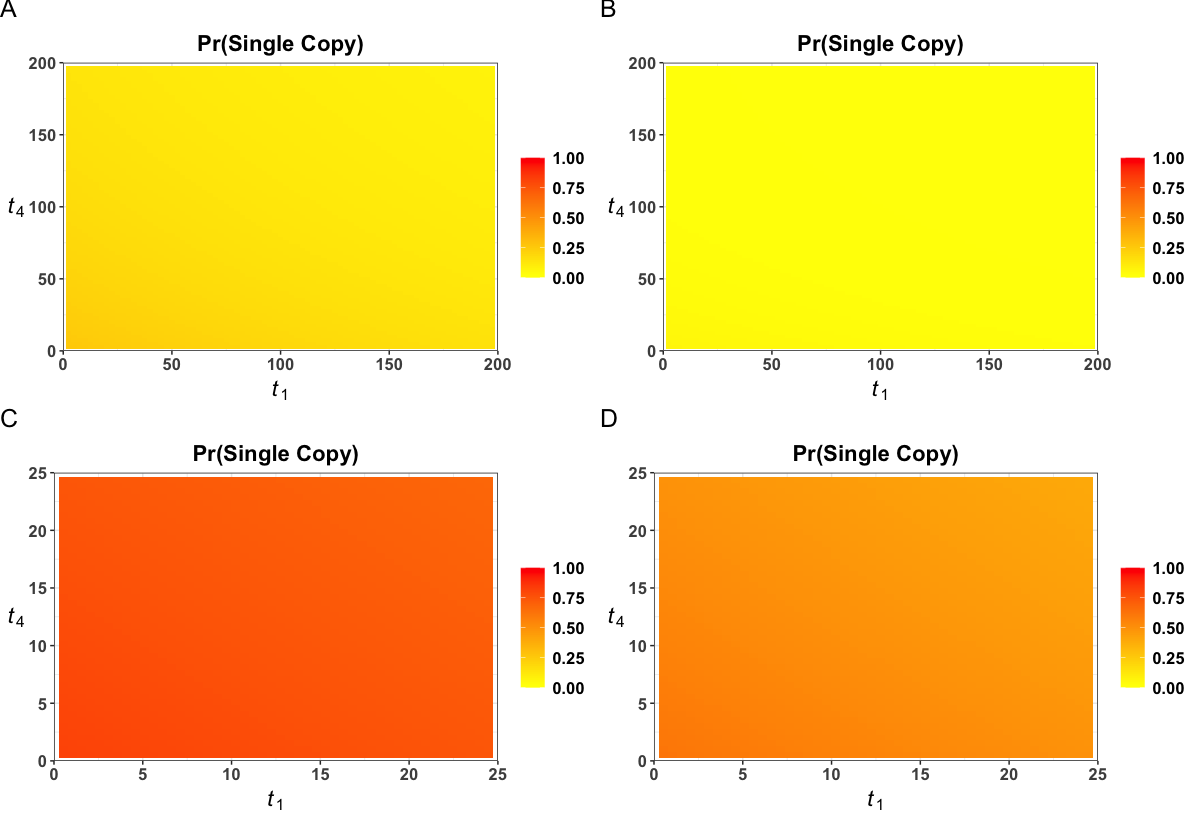
Fig S7. Probabilities of single-copy genes. A) The probability of single-copy genes with reasonable rates of duplication and loss (λ=0.002, μ=0.002). Here, branch length *t_3_* = 198.9. Branch length *t_1_* varies from 0.0001 to 200, while branch length *t_4_* varies from 0.0001 to 198.8; branch length *t_2_* is constrained such that the sum of *t_2_* and *t_4_* equals *t_3_*. B) The probability of single-copy genes with high rates of duplication and loss (λ=0.005, μ=0.005). Same branch lengths as in panel A. C) The probability of single-copy genes with reasonable rates of duplication and loss (λ=0.002, μ=0.002). Here, branch length *t_3_* = 25. Branch length *t_1_* varies from 0.0001 to 25, while branch length *t_4_* varies from 0.0001 to 24.9; branch length *t_2_* is constrained such that the sum of *t_2_* and *t_4_* equals *t_3_*. D) The probability of single copy genes with high rates of duplication and loss (λ=0.005, μ=0.005). Same branch lengths as in panel C.

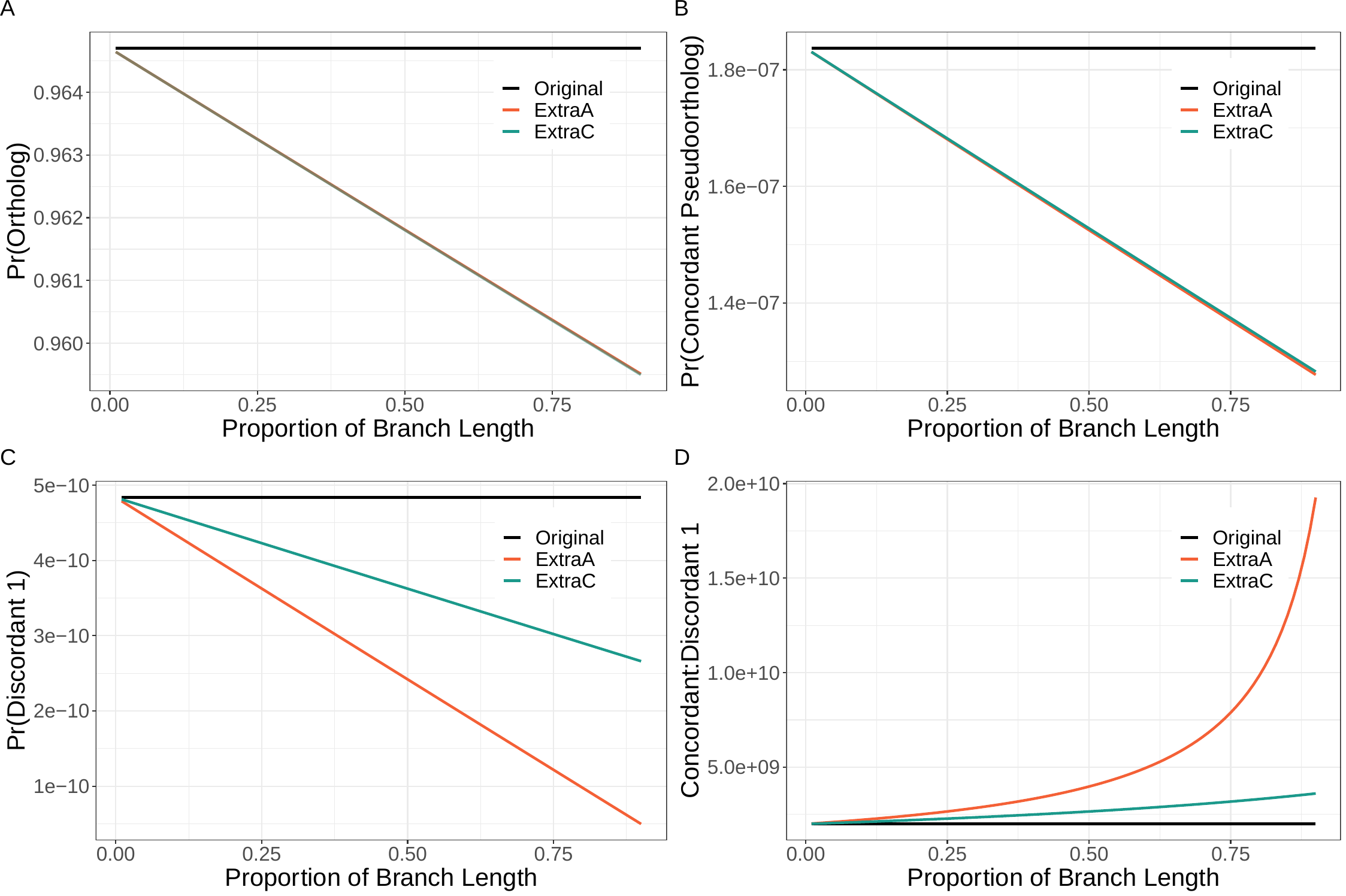
Fig S8. Results on larger trees. The original scenario includes a single branch per species. Here ExtraA includes an extra branch sister to Species A and ExtraC includes an extra branch sister to species C. The parameters for all panels are: μ = 0.004, λ=0.002, *t_1_*=1, *t_2_*=1, *t_3_*=2, *t_4_*/*t_5_*=1. The length of the added branch is proportional to branch *t_4_*. A) The probability of orthologs. B) The probability of concordant pseudoorthologs. C) The probability of discordant pseudoortholog 1. D) The ratio of the concordant topologies to the probability of discordant pseudoortholog 1.

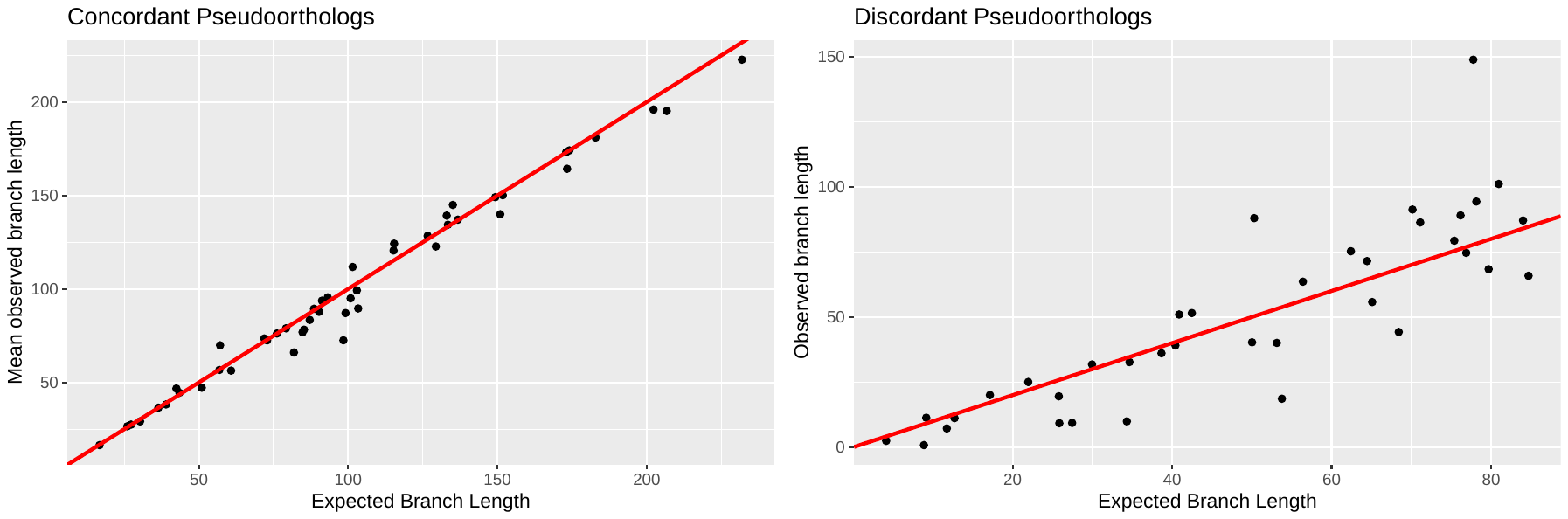
Fig S9. Results of simulations to evaluate branch length predictions. Red lines indicate the predicted 1:1 relationship between the expected internal branch lengths and the observed internal branch lengths in simulated datasets. Each datapoint is the average internal branch length of all simulations under a set of parameters that match the relevant scenario. For each set of parameters, we conducted 10,000 simulations. A) The expected versus observed internal branch lengths of concordant pseudoortholog 1 (Figure 1d; Figure S2a). B) The expected versus observed internal branch lengths of discordant pseudoortholog 1 when there were two copies at the end of branch *t_1_* (Figure 1e; Figure S3a).
Table S1. The parameters that maximize the relative probabilities of pseudoorthologs, maximize the relative probability of discordant pseudoorthologs, and minimize the ratio of concordant to discordant topologies. Discordant probabilities and ratios refer only to discordant pseudoortholog 1, but the probabilities of the two discordant pseudoorthologs are equal.

|  | **Pseudoorthologs** | **Discordant** | **Concordant: Discordant** |
| --- | --- | --- | --- |
| μ | 0.0049 | 0.0050 | 0.0050 |
| λ | 0.0050 | 0.0050 | 0.0049 |
| t_1_ | 199.8807 | 199.9939 | 193.4689 |
| t_2_ | 0.5893 | 1.1028 | 0.2272 |
| t_3_ | 199.8666 | 199.4551 | 199.8339 |
| t_4_ | 199.2773 | 198.3523 | 199.6066 |
| t_5_ | 199.2773 | 198.3523 | 199.6066 |
|  | Pr(Pseudoorthologs) = 0.285  Pr(Orthologs) = 0.715 | Pr(Discordant Pseudo) = 0.095  Pr(Concordant Pseudo) = 0.099  Pr(Orthologs) = 0.711 | Concordant: Discordant = 8.53  Pr(Discordant Pseudo) = 0.095  Pr(Concordant Pseudo) = 0.096  Pr(Orthologs) = 0.714 |

Table S2. The parameters that maximize the absolute probabilities of pseudoorthologs and the probability of discordant pseudoorthologs.

|  | **Pseudoorthologs** | **Discordant** |
| --- | --- | --- |
| μ | 0.0049 | 0.0043 |
| λ | 0.0031 | 0.0024 |
| t_1_ | 136.5054 | 152.1832 |
| t_2_ | 135.8794 | 0.1056 |
| t_3_ | 136.7011 | 150.6176 |
| t_4_ | 0.8217329 | 150.512 |
| t_5_ | 0.8217329 | 150.512 |
|  | Pr(Pseudoorthologs) = 0.0082  Pr(Orthologs) = 0.0779 | Pr(Discordant) = 0.0029  Pr(Concordant) = 0.0014  Pr(Orthologs) = 0.0362 |

Table S3. Bounds on parameters used when performing optimization.

|  | **Minimum** | **Maximum** |
| --- | --- | --- |
| μ | 0.0001 | 0.005 |
| λ | 0.0001 | 0.005 |
| t_1_ | 0.0001 | 200 |
| t_2_ | 0.0001 | 200 |
| t_3_ | 0.0001 | 200 |

Table S4. Priors used in SimPhy simulations.

|  | **Minimum** | **Maximum** |
| --- | --- | --- |
| μ | 0.0001 | 0.005 |
| λ | 0.0001 | 0.005 |
| t_1_ | 0.01 | 200 |
| t_2_ | 0.01 | 200 |
| t_3_ | 0.01 | 200 |
