## Appendix A for "The frequency and topology of pseudoorthologs"

### Appendix A: Probabilities of Pseudoorthologs

#### Calculate transition probabilities

The code below calculates the probability of a transition from a=1 copies to n copies given a duplication rate (lambda), a loss rate (mu), and a branch length (t).

```
birthdeathprob <- function(a,n,lambda,mu, t)
{
  if (n>0 & isTRUE(all.equal(mu, lambda)))
  {
    prob=((lambda*t)^(n-1))/((1+lambda*t)^(n+1))
  }
  else if (n==0 & isTRUE(all.equal(mu, lambda)))
  {
    prob=(lambda*t)/(1+lambda*t)
  }

  else if (n > 0){
    alpha = (mu * (exp((lambda - mu) * t) - 1)) / (lambda * exp((lambda - mu) * t) - mu)
    beta =  (lambda * (exp((lambda - mu) * t) - 1)) / (lambda * exp((lambda - mu) * t) - mu)

    prob = (1-alpha)*(1-beta)*beta^(n-1)
  }
  else if (n==0){
    alpha = (mu * (exp((lambda - mu) * t) - 1)) / (lambda * exp((lambda - mu) * t) - mu)

    prob = alpha
  }

  return(prob)
}
```

#### Calculate ortholog probabilities

The following code computes the probability of orthologs by adding probabilities of different ortholog configurations from Supporting Figure S1.

```
orthologprob <- function(lambda, mu, t1, t2, t3, t4, t5)
{

  # Figure S1A: True ortholog 1- no transitions
  prortholog <- birthdeathprob(a=1, n=1, lambda=lambda, mu=mu, t=t1) *
    birthdeathprob(a=1, n=1, lambda=lambda, mu=mu, t=t2) *
    birthdeathprob(a=1, n=1, lambda=lambda, mu=mu, t=t3) *
    birthdeathprob(a=1, n=1, lambda=lambda, mu=mu, t=t4) *
```

```
birthdeathprob(a=1, n=1, lambda=lambda, mu=mu, t=t5) +

# Figure S1B: True ortholog 2 - two copies
birthdeathprob(a=1, n=2, lambda=lambda, mu=mu, t=t1) * # 1 -> 2 on branch t1
birthdeathprob(a=1, n=1, lambda=lambda, mu=mu, t=t2) * # tree 1
birthdeathprob(a=1, n=1, lambda=lambda, mu=mu, t=t3) * # tree 1
birthdeathprob(a=1, n=1, lambda=lambda, mu=mu, t=t4) * # tree 1
birthdeathprob(a=1, n=1, lambda=lambda, mu=mu, t=t5) * # tree 1
birthdeathprob(a=1, n=0, lambda=lambda, mu=mu, t=t2) * # tree 2
birthdeathprob(a=1, n=0, lambda=lambda, mu=mu, t=t3) * # tree 2
2 +# two arrangements

# Figure S1C: True ortholog 3 - two copies
birthdeathprob(a=1, n=2, lambda=lambda, mu=mu, t=t1) * # 1 -> 2 on branch t1
birthdeathprob(a=1, n=1, lambda=lambda, mu=mu, t=t2) * # tree 1
birthdeathprob(a=1, n=1, lambda=lambda, mu=mu, t=t3) * # tree 1
birthdeathprob(a=1, n=1, lambda=lambda, mu=mu, t=t4) * # tree 1
birthdeathprob(a=1, n=1, lambda=lambda, mu=mu, t=t5) * # tree 1
birthdeathprob(a=1, n=1, lambda=lambda, mu=mu, t=t2) * # tree 1
birthdeathprob(a=1, n=0, lambda=lambda, mu=mu, t=t3) * # tree 0
birthdeathprob(a=1, n=0, lambda=lambda, mu=mu, t=t4) * # tree 0
birthdeathprob(a=1, n=0, lambda=lambda, mu=mu, t=t5) * # tree 0
2 +# two arrangements

# Figure S1D: True ortholog 4 - three copies
birthdeathprob(a=1, n=3, lambda=lambda, mu=mu, t=t1) * # 1 -> 3 on branch t1
birthdeathprob(a=1, n=1, lambda=lambda, mu=mu, t=t2) * # tree 1
birthdeathprob(a=1, n=1, lambda=lambda, mu=mu, t=t3) * # tree 1
birthdeathprob(a=1, n=1, lambda=lambda, mu=mu, t=t4) * # tree 1
birthdeathprob(a=1, n=1, lambda=lambda, mu=mu, t=t5) * # tree 1
birthdeathprob(a=1, n=0, lambda=lambda, mu=mu, t=t2) * # tree 2
birthdeathprob(a=1, n=0, lambda=lambda, mu=mu, t=t3) * # tree 2
birthdeathprob(a=1, n=0, lambda=lambda, mu=mu, t=t2) * # tree 3
birthdeathprob(a=1, n=0, lambda=lambda, mu=mu, t=t3) * # tree 3
3 +# three arrangements


# Figure S1E: True ortholog 5 - three copies
birthdeathprob(a=1, n=3, lambda=lambda, mu=mu, t=t1) * # 1 -> 3 on branch t1
birthdeathprob(a=1, n=1, lambda=lambda, mu=mu, t=t2) * # tree 1
birthdeathprob(a=1, n=1, lambda=lambda, mu=mu, t=t3) * # tree 1
birthdeathprob(a=1, n=1, lambda=lambda, mu=mu, t=t4) * # tree 1
birthdeathprob(a=1, n=1, lambda=lambda, mu=mu, t=t5) * # tree 1
birthdeathprob(a=1, n=1, lambda=lambda, mu=mu, t=t2) * # tree 2
birthdeathprob(a=1, n=0, lambda=lambda, mu=mu, t=t3) * # tree 2
birthdeathprob(a=1, n=0, lambda=lambda, mu=mu, t=t4) * # tree 2
birthdeathprob(a=1, n=0, lambda=lambda, mu=mu, t=t5) * # tree 2
birthdeathprob(a=1, n=0, lambda=lambda, mu=mu, t=t2) * # tree 3
birthdeathprob(a=1, n=0, lambda=lambda, mu=mu, t=t3) * # tree 3
6 +# six arrangements

# Figure S1F: True ortholog 6 - three copies
birthdeathprob(a=1, n=3, lambda=lambda, mu=mu, t=t1) * # 1 -> 3 on branch t1
```

```
    birthdeathprob(a=1, n=1, lambda=lambda, mu=mu, t=t2) * # tree 2
    birthdeathprob(a=1, n=1, lambda=lambda, mu=mu, t=t3) * # tree 2
    birthdeathprob(a=1, n=1, lambda=lambda, mu=mu, t=t4) * # tree 2
    birthdeathprob(a=1, n=1, lambda=lambda, mu=mu, t=t5) * # tree 2
    birthdeathprob(a=1, n=1, lambda=lambda, mu=mu, t=t2) * # tree 2
    birthdeathprob(a=1, n=0, lambda=lambda, mu=mu, t=t3) * # tree 2
    birthdeathprob(a=1, n=0, lambda=lambda, mu=mu, t=t4) * # tree 2
    birthdeathprob(a=1, n=0, lambda=lambda, mu=mu, t=t5) * # tree 2
    birthdeathprob(a=1, n=1, lambda=lambda, mu=mu, t=t2) * # tree 3
    birthdeathprob(a=1, n=0, lambda=lambda, mu=mu, t=t3) * # tree 3
    birthdeathprob(a=1, n=0, lambda=lambda, mu=mu, t=t4) * # tree 3
    birthdeathprob(a=1, n=0, lambda=lambda, mu=mu, t=t5) * # tree 3
    3 # three arrangements

    return(prortholog)

}
```

#### Calculate concordant pseudoortholog probabilities

The following code calculates the probability of the concordant pseudoortholog. Configurations are shown in Supporting Figure S2.

```
pseudoconcordprob <- function(lambda, mu, t1, t2, t3, t4, t5)
{

  # Figure S2A: Concordant pseudoortholog 1 - two copies
  pseudoconcord <- birthdeathprob(a=1, n=2, lambda=lambda, mu=mu, t=t1) * # 1 -> 2 on branch t1
    birthdeathprob(a=1, n=1, lambda=lambda, mu=mu, t=t2) * # tree 1
    birthdeathprob(a=1, n=0, lambda=lambda, mu=mu, t=t3) * # tree 1
    birthdeathprob(a=1, n=1, lambda=lambda, mu=mu, t=t4) * # tree 1
    birthdeathprob(a=1, n=1, lambda=lambda, mu=mu, t=t5) * # tree 1
    birthdeathprob(a=1, n=0, lambda=lambda, mu=mu, t=t2) * # tree 2
    birthdeathprob(a=1, n=1, lambda=lambda, mu=mu, t=t3) * # tree 2
    2 +# two arrangements

    # Figure S2B: Concordant pseudoortholog 2 - two copies
    birthdeathprob(a=1, n=2, lambda=lambda, mu=mu, t=t1) * # 1 -> 2 on branch t1
    birthdeathprob(a=1, n=1, lambda=lambda, mu=mu, t=t2) * # tree 1
    birthdeathprob(a=1, n=0, lambda=lambda, mu=mu, t=t3) * # tree 1
    birthdeathprob(a=1, n=1, lambda=lambda, mu=mu, t=t4) * # tree 1
    birthdeathprob(a=1, n=1, lambda=lambda, mu=mu, t=t5) * # tree 1
    birthdeathprob(a=1, n=1, lambda=lambda, mu=mu, t=t2) * # tree 2
    birthdeathprob(a=1, n=1, lambda=lambda, mu=mu, t=t3) * # tree 2
    birthdeathprob(a=1, n=0, lambda=lambda, mu=mu, t=t4) * # tree 2
    birthdeathprob(a=1, n=0, lambda=lambda, mu=mu, t=t5) * # tree 2
    2 +# two arrangements

    # Figure S2D: Concordant pseudoortholog 3 - three copies
    birthdeathprob(a=1, n=3, lambda=lambda, mu=mu, t=t1) * # 1 -> 3 on branch t1
    birthdeathprob(a=1, n=0, lambda=lambda, mu=mu, t=t2) * # tree 1
    birthdeathprob(a=1, n=0, lambda=lambda, mu=mu, t=t3) * # tree 1
```

```
birthdeathprob(a=1, n=1, lambda=lambda, mu=mu, t=t2) * # tree 2
birthdeathprob(a=1, n=0, lambda=lambda, mu=mu, t=t3) * # tree 2
birthdeathprob(a=1, n=1, lambda=lambda, mu=mu, t=t4) * # tree 2
birthdeathprob(a=1, n=1, lambda=lambda, mu=mu, t=t5) * # tree 2
birthdeathprob(a=1, n=0, lambda=lambda, mu=mu, t=t2) * # tree 3
birthdeathprob(a=1, n=1, lambda=lambda, mu=mu, t=t3) * # tree 3
6 + # six arrangements

# S2E: Concordant pseudoortholog 4 - three copies
birthdeathprob(a=1, n=3, lambda=lambda, mu=mu, t=t1) * # 1 -> 3 on branch t1
birthdeathprob(a=1, n=0, lambda=lambda, mu=mu, t=t2) * # tree 1
birthdeathprob(a=1, n=0, lambda=lambda, mu=mu, t=t3) * # tree 1
birthdeathprob(a=1, n=1, lambda=lambda, mu=mu, t=t2) * # tree 3
birthdeathprob(a=1, n=0, lambda=lambda, mu=mu, t=t3) * # tree 3
birthdeathprob(a=1, n=1, lambda=lambda, mu=mu, t=t4) * # tree 3
birthdeathprob(a=1, n=1, lambda=lambda, mu=mu, t=t5) * # tree 3
birthdeathprob(a=1, n=1, lambda=lambda, mu=mu, t=t2) * # tree 2
birthdeathprob(a=1, n=1, lambda=lambda, mu=mu, t=t3) * # tree 2
birthdeathprob(a=1, n=0, lambda=lambda, mu=mu, t=t4) * # tree 2
birthdeathprob(a=1, n=0, lambda=lambda, mu=mu, t=t5) * # tree 2
6 + # six arrangements

# S2F: Concordant pseudoortholog 5 - three copies
birthdeathprob(a=1, n=3, lambda=lambda, mu=mu, t=t1) * # 1 -> 3 on branch t1
birthdeathprob(a=1, n=1, lambda=lambda, mu=mu, t=t2) * # tree 1
birthdeathprob(a=1, n=0, lambda=lambda, mu=mu, t=t3) * # tree 1
birthdeathprob(a=1, n=0, lambda=lambda, mu=mu, t=t4) * # tree 1
birthdeathprob(a=1, n=0, lambda=lambda, mu=mu, t=t5) * # tree 1
birthdeathprob(a=1, n=1, lambda=lambda, mu=mu, t=t2) * # tree 3
birthdeathprob(a=1, n=0, lambda=lambda, mu=mu, t=t3) * # tree 3
birthdeathprob(a=1, n=1, lambda=lambda, mu=mu, t=t4) * # tree 3
birthdeathprob(a=1, n=1, lambda=lambda, mu=mu, t=t5) * # tree 3
birthdeathprob(a=1, n=0, lambda=lambda, mu=mu, t=t2) * # tree 2
birthdeathprob(a=1, n=1, lambda=lambda, mu=mu, t=t3) * # tree 2
6 + # six arrangements


# S2G: Concordant pseudoortholog 6 - three copies
birthdeathprob(a=1, n=3, lambda=lambda, mu=mu, t=t1) * # 1 -> 3 on branch t1
birthdeathprob(a=1, n=1, lambda=lambda, mu=mu, t=t2) * # tree 1
birthdeathprob(a=1, n=0, lambda=lambda, mu=mu, t=t3) * # tree 1
birthdeathprob(a=1, n=0, lambda=lambda, mu=mu, t=t4) * # tree 1
birthdeathprob(a=1, n=0, lambda=lambda, mu=mu, t=t5) * # tree 1
birthdeathprob(a=1, n=1, lambda=lambda, mu=mu, t=t2) * # tree 2
birthdeathprob(a=1, n=0, lambda=lambda, mu=mu, t=t3) * # tree 2
birthdeathprob(a=1, n=1, lambda=lambda, mu=mu, t=t4) * # tree 2
birthdeathprob(a=1, n=1, lambda=lambda, mu=mu, t=t5) * # tree 2
birthdeathprob(a=1, n=1, lambda=lambda, mu=mu, t=t2) * # tree 3
birthdeathprob(a=1, n=1, lambda=lambda, mu=mu, t=t3) * # tree 3
birthdeathprob(a=1, n=0, lambda=lambda, mu=mu, t=t4) * # tree 3
birthdeathprob(a=1, n=0, lambda=lambda, mu=mu, t=t5) * # tree 3
6 + # six arrangements
```

```
    # S2H: Concordant pseudoortholog 7 - three copies
    birthdeathprob(a=1, n=3, lambda=lambda, mu=mu, t=t1) * # 1 -> 3 on branch t1
    birthdeathprob(a=1, n=1, lambda=lambda, mu=mu, t=t2) * # tree 1
    birthdeathprob(a=1, n=0, lambda=lambda, mu=mu, t=t3) * # tree 1
    birthdeathprob(a=1, n=1, lambda=lambda, mu=mu, t=t4) * # tree 1
    birthdeathprob(a=1, n=0, lambda=lambda, mu=mu, t=t5) * # tree 1
    birthdeathprob(a=1, n=1, lambda=lambda, mu=mu, t=t2) * # tree 2
    birthdeathprob(a=1, n=0, lambda=lambda, mu=mu, t=t3) * # tree 2
    birthdeathprob(a=1, n=0, lambda=lambda, mu=mu, t=t4) * # tree 2
    birthdeathprob(a=1, n=1, lambda=lambda, mu=mu, t=t5) * # tree 2
    birthdeathprob(a=1, n=0, lambda=lambda, mu=mu, t=t2) * # tree 3
    birthdeathprob(a=1, n=1, lambda=lambda, mu=mu, t=t3) * # tree 3
    6* # six arrangements
    (1/3) + # because the topology is only concordant 1/3 of the time

    # S2I: Concordant pseudoortholog 8 - three copies
    birthdeathprob(a=1, n=3, lambda=lambda, mu=mu, t=t1) * # 1 -> 3 on branch t1
    birthdeathprob(a=1, n=1, lambda=lambda, mu=mu, t=t2) * # tree 1
    birthdeathprob(a=1, n=0, lambda=lambda, mu=mu, t=t3) * # tree 1
    birthdeathprob(a=1, n=1, lambda=lambda, mu=mu, t=t4) * # tree 1
    birthdeathprob(a=1, n=0, lambda=lambda, mu=mu, t=t5) * # tree 1
    birthdeathprob(a=1, n=1, lambda=lambda, mu=mu, t=t2) * # tree 3
    birthdeathprob(a=1, n=0, lambda=lambda, mu=mu, t=t3) * # tree 3
    birthdeathprob(a=1, n=0, lambda=lambda, mu=mu, t=t4) * # tree 3
    birthdeathprob(a=1, n=1, lambda=lambda, mu=mu, t=t5) * # tree 3
    birthdeathprob(a=1, n=1, lambda=lambda, mu=mu, t=t2) * # tree 2
    birthdeathprob(a=1, n=1, lambda=lambda, mu=mu, t=t3) * # tree 2
    birthdeathprob(a=1, n=0, lambda=lambda, mu=mu, t=t4) * # tree 2
    birthdeathprob(a=1, n=0, lambda=lambda, mu=mu, t=t5) * # tree 2
    6 * # six arrangements
    (1/3) +# because the topology is only concordant 1/3 of the time

    # S2C: Concordant pseudoortholog 9 - two lineage duplicate
    birthdeathprob(a=1, n=1, lambda=lambda, mu=mu, t=t1) * # tree 1
    birthdeathprob(a=1, n=2, lambda=lambda, mu=mu, t=t2) * # tree 1
    birthdeathprob(a=1, n=1, lambda=lambda, mu=mu, t=t3) * # tree 1
    birthdeathprob(a=1, n=1, lambda=lambda, mu=mu, t=t4) * # tree 1
    birthdeathprob(a=1, n=0, lambda=lambda, mu=mu, t=t5) * # tree 1
    birthdeathprob(a=1, n=0, lambda=lambda, mu=mu, t=t4) * # tree 2
    birthdeathprob(a=1, n=1, lambda=lambda, mu=mu, t=t5) * # tree 2
    2  # two arrangements

    return(pseudoconcord)

}
```

#### Calculate discordant pseudoortholog probabilities

The following code calculates the probability of discordant pseudoortholog 1. Configurations are shown in Supporting Figure S4.

```
pseudodiscord1prob <- function(lambda, mu, t1, t2, t3, t4, t5)
{
  # Figure S3A: Discordant pseudoortholog 1 - 1 - two copies
  pseudo1discord1 <- birthdeathprob(a=1, n=2, lambda=lambda, mu=mu, t=t1) * # 1 -> 2 on branch t1
    birthdeathprob(a=1, n=1, lambda=lambda, mu=mu, t=t2) * # tree 1
    birthdeathprob(a=1, n=0, lambda=lambda, mu=mu, t=t3) * # tree 1
    birthdeathprob(a=1, n=1, lambda=lambda, mu=mu, t=t4) * # tree 1
    birthdeathprob(a=1, n=0, lambda=lambda, mu=mu, t=t5) * # tree 1
    birthdeathprob(a=1, n=1, lambda=lambda, mu=mu, t=t2) * # tree 2
    birthdeathprob(a=1, n=1, lambda=lambda, mu=mu, t=t3) * # tree 2
    birthdeathprob(a=1, n=0, lambda=lambda, mu=mu, t=t4) * # tree 2
    birthdeathprob(a=1, n=1, lambda=lambda, mu=mu, t=t5) * # tree 2
    2 +# two arrangements

    # Figure S3C: Discordant pseudoortholog 1 - 2 - three copies
    birthdeathprob(a=1, n=3, lambda=lambda, mu=mu, t=t1) * # 1 -> 3 on branch t1
    birthdeathprob(a=1, n=0, lambda=lambda, mu=mu, t=t2) * # tree 1
    birthdeathprob(a=1, n=0, lambda=lambda, mu=mu, t=t3) * # tree 1
    birthdeathprob(a=1, n=1, lambda=lambda, mu=mu, t=t2) * # tree 2
    birthdeathprob(a=1, n=0, lambda=lambda, mu=mu, t=t3) * # tree 2
    birthdeathprob(a=1, n=1, lambda=lambda, mu=mu, t=t4) * # tree 2
    birthdeathprob(a=1, n=0, lambda=lambda, mu=mu, t=t5) * # tree 2
    birthdeathprob(a=1, n=1, lambda=lambda, mu=mu, t=t2) * # tree 3
    birthdeathprob(a=1, n=1, lambda=lambda, mu=mu, t=t3) * # tree 3
    birthdeathprob(a=1, n=0, lambda=lambda, mu=mu, t=t4) * # tree 3
    birthdeathprob(a=1, n=1, lambda=lambda, mu=mu, t=t5) * # tree 3
    6 + # six arrangements

    # Figure S3E: Discordant pseudoortholog 1 - 3 - three copies
    birthdeathprob(a=1, n=3, lambda=lambda, mu=mu, t=t1) * # 1 -> 3 on branch t1
    birthdeathprob(a=1, n=1, lambda=lambda, mu=mu, t=t2) * # tree 1
    birthdeathprob(a=1, n=0, lambda=lambda, mu=mu, t=t3) * # tree 1
    birthdeathprob(a=1, n=0, lambda=lambda, mu=mu, t=t4) * # tree 1
    birthdeathprob(a=1, n=0, lambda=lambda, mu=mu, t=t5) * # tree 1
    birthdeathprob(a=1, n=1, lambda=lambda, mu=mu, t=t2) * # tree 2
    birthdeathprob(a=1, n=0, lambda=lambda, mu=mu, t=t3) * # tree 2
    birthdeathprob(a=1, n=1, lambda=lambda, mu=mu, t=t4) * # tree 2
    birthdeathprob(a=1, n=0, lambda=lambda, mu=mu, t=t5) * # tree 2
    birthdeathprob(a=1, n=1, lambda=lambda, mu=mu, t=t2) * # tree 3
    birthdeathprob(a=1, n=1, lambda=lambda, mu=mu, t=t3) * # tree 3
    birthdeathprob(a=1, n=0, lambda=lambda, mu=mu, t=t4) * # tree 3
    birthdeathprob(a=1, n=1, lambda=lambda, mu=mu, t=t5) * # tree 3
    6 + # six arrangements

    # Figure S3G: Discordant pseudoortholog 1 - 4 - three copies
    birthdeathprob(a=1, n=3, lambda=lambda, mu=mu, t=t1) * # 1 -> 3 on branch t1
    birthdeathprob(a=1, n=0, lambda=lambda, mu=mu, t=t2) * # tree 1
    birthdeathprob(a=1, n=1, lambda=lambda, mu=mu, t=t3) * # tree 1
    birthdeathprob(a=1, n=1, lambda=lambda, mu=mu, t=t2) * # tree 2
    birthdeathprob(a=1, n=0, lambda=lambda, mu=mu, t=t3) * # tree 2
    birthdeathprob(a=1, n=0, lambda=lambda, mu=mu, t=t4) * # tree 2
    birthdeathprob(a=1, n=1, lambda=lambda, mu=mu, t=t5) * # tree 2
    birthdeathprob(a=1, n=1, lambda=lambda, mu=mu, t=t2) * # tree 3
```

```
    birthdeathprob(a=1, n=0, lambda=lambda, mu=mu, t=t3) * # tree 3
    birthdeathprob(a=1, n=1, lambda=lambda, mu=mu, t=t4) * # tree 3
    birthdeathprob(a=1, n=0, lambda=lambda, mu=mu, t=t5) * # tree 3
    6* # six arrangements
    (1/3) + # because the topology is only concordant 1/3 of the time

    # Figure S3I: Discordant pseudoortholog 1 - 5 - three copies
    birthdeathprob(a=1, n=3, lambda=lambda, mu=mu, t=t1) * # 1 -> 3 on branch t1
    birthdeathprob(a=1, n=1, lambda=lambda, mu=mu, t=t2) * # tree 1
    birthdeathprob(a=1, n=1, lambda=lambda, mu=mu, t=t3) * # tree 1
    birthdeathprob(a=1, n=0, lambda=lambda, mu=mu, t=t4) * # tree 1
    birthdeathprob(a=1, n=0, lambda=lambda, mu=mu, t=t5) * # tree 1
    birthdeathprob(a=1, n=1, lambda=lambda, mu=mu, t=t2) * # tree 2
    birthdeathprob(a=1, n=0, lambda=lambda, mu=mu, t=t3) * # tree 2
    birthdeathprob(a=1, n=0, lambda=lambda, mu=mu, t=t4) * # tree 2
    birthdeathprob(a=1, n=1, lambda=lambda, mu=mu, t=t5) * # tree 2
    birthdeathprob(a=1, n=1, lambda=lambda, mu=mu, t=t2) * # tree 3
    birthdeathprob(a=1, n=0, lambda=lambda, mu=mu, t=t3) * # tree 3
    birthdeathprob(a=1, n=1, lambda=lambda, mu=mu, t=t4) * # tree 3
    birthdeathprob(a=1, n=0, lambda=lambda, mu=mu, t=t5) * # tree 3
    6 * # six arrangements
    (1/3) # because the topology is only concordant 1/3 of the time

    return(pseudo1discord1)
}
```

The following code calculates the probability of discordant pseudoortholog 2. Configurations are shown in Supporting Figure S4.

```
pseudodiscord2prob <- function(lambda, mu, t1, t2, t3, t4, t5)
{
  # Figure S3B: Discordant pseudoortholog 2 - 1 - two copies
  pseudo1discord2 <- birthdeathprob(a=1, n=2, lambda=lambda, mu=mu, t=t1) * # 1 -> 2 on branch t1
    birthdeathprob(a=1, n=1, lambda=lambda, mu=mu, t=t2) * # tree 1
    birthdeathprob(a=1, n=0, lambda=lambda, mu=mu, t=t3) * # tree 1
    birthdeathprob(a=1, n=0, lambda=lambda, mu=mu, t=t4) * # tree 1
    birthdeathprob(a=1, n=1, lambda=lambda, mu=mu, t=t5) * # tree 1
    birthdeathprob(a=1, n=1, lambda=lambda, mu=mu, t=t2) * # tree 2
    birthdeathprob(a=1, n=1, lambda=lambda, mu=mu, t=t3) * # tree 2
    birthdeathprob(a=1, n=1, lambda=lambda, mu=mu, t=t4) * # tree 2
    birthdeathprob(a=1, n=0, lambda=lambda, mu=mu, t=t5) * # tree 2
    2 +# two arrangements

    # Figure S3D: Discordant pseudoortholog 2 - 2 - three copies
    birthdeathprob(a=1, n=3, lambda=lambda, mu=mu, t=t1) * # 1 -> 3 on branch t1
    birthdeathprob(a=1, n=0, lambda=lambda, mu=mu, t=t2) * # tree 1
    birthdeathprob(a=1, n=0, lambda=lambda, mu=mu, t=t3) * # tree 1
    birthdeathprob(a=1, n=1, lambda=lambda, mu=mu, t=t2) * # tree 2
    birthdeathprob(a=1, n=0, lambda=lambda, mu=mu, t=t3) * # tree 2
    birthdeathprob(a=1, n=0, lambda=lambda, mu=mu, t=t4) * # tree 2
    birthdeathprob(a=1, n=1, lambda=lambda, mu=mu, t=t5) * # tree 2
    birthdeathprob(a=1, n=1, lambda=lambda, mu=mu, t=t2) * # tree 3
    birthdeathprob(a=1, n=1, lambda=lambda, mu=mu, t=t3) * # tree 3
```

```
  birthdeathprob(a=1, n=1, lambda=lambda, mu=mu, t=t4) * # tree 3
  birthdeathprob(a=1, n=0, lambda=lambda, mu=mu, t=t5) * # tree 3
  6 + # six arrangements

  # Figure S3F: Discordant pseudoortholog 2 - 3 - three copies
  birthdeathprob(a=1, n=3, lambda=lambda, mu=mu, t=t1) * # 1 -> 3 on branch t1
  birthdeathprob(a=1, n=1, lambda=lambda, mu=mu, t=t2) * # tree 1
  birthdeathprob(a=1, n=0, lambda=lambda, mu=mu, t=t3) * # tree 1
  birthdeathprob(a=1, n=0, lambda=lambda, mu=mu, t=t4) * # tree 1
  birthdeathprob(a=1, n=0, lambda=lambda, mu=mu, t=t5) * # tree 1
  birthdeathprob(a=1, n=1, lambda=lambda, mu=mu, t=t2) * # tree 2
  birthdeathprob(a=1, n=0, lambda=lambda, mu=mu, t=t3) * # tree 2
  birthdeathprob(a=1, n=0, lambda=lambda, mu=mu, t=t4) * # tree 2
  birthdeathprob(a=1, n=1, lambda=lambda, mu=mu, t=t5) * # tree 2
  birthdeathprob(a=1, n=1, lambda=lambda, mu=mu, t=t2) * # tree 3
  birthdeathprob(a=1, n=1, lambda=lambda, mu=mu, t=t3) * # tree 3
  birthdeathprob(a=1, n=1, lambda=lambda, mu=mu, t=t4) * # tree 3
  birthdeathprob(a=1, n=0, lambda=lambda, mu=mu, t=t5) * # tree 3
  6 + # six arrangements

  # Figure S3H: Discordant pseudoortholog 2 - 4 - three copies
  birthdeathprob(a=1, n=3, lambda=lambda, mu=mu, t=t1) * # 1 -> 3 on branch t1
  birthdeathprob(a=1, n=0, lambda=lambda, mu=mu, t=t2) * # tree 1
  birthdeathprob(a=1, n=1, lambda=lambda, mu=mu, t=t3) * # tree 1
  birthdeathprob(a=1, n=1, lambda=lambda, mu=mu, t=t2) * # tree 2
  birthdeathprob(a=1, n=0, lambda=lambda, mu=mu, t=t3) * # tree 2
  birthdeathprob(a=1, n=1, lambda=lambda, mu=mu, t=t4) * # tree 2
  birthdeathprob(a=1, n=0, lambda=lambda, mu=mu, t=t5) * # tree 2
  birthdeathprob(a=1, n=1, lambda=lambda, mu=mu, t=t2) * # tree 3
  birthdeathprob(a=1, n=0, lambda=lambda, mu=mu, t=t3) * # tree 3
  birthdeathprob(a=1, n=0, lambda=lambda, mu=mu, t=t4) * # tree 3
  birthdeathprob(a=1, n=1, lambda=lambda, mu=mu, t=t5) * # tree 3
  6* # six arrangements
  (1/3) + # because the topology is only concordant 1/3 of the time

  # Figure S3J: Discordant pseudoortholog 2 - 5 - three copies
  birthdeathprob(a=1, n=3, lambda=lambda, mu=mu, t=t1) * # 1 -> 3 on branch t1
  birthdeathprob(a=1, n=1, lambda=lambda, mu=mu, t=t2) * # tree 1
  birthdeathprob(a=1, n=0, lambda=lambda, mu=mu, t=t3) * # tree 1
  birthdeathprob(a=1, n=1, lambda=lambda, mu=mu, t=t4) * # tree 1
  birthdeathprob(a=1, n=0, lambda=lambda, mu=mu, t=t5) * # tree 1
  birthdeathprob(a=1, n=1, lambda=lambda, mu=mu, t=t2) * # tree 2
  birthdeathprob(a=1, n=1, lambda=lambda, mu=mu, t=t3) * # tree 2
  birthdeathprob(a=1, n=0, lambda=lambda, mu=mu, t=t4) * # tree 2
  birthdeathprob(a=1, n=0, lambda=lambda, mu=mu, t=t5) * # tree 2
  birthdeathprob(a=1, n=1, lambda=lambda, mu=mu, t=t2) * # tree 3
  birthdeathprob(a=1, n=0, lambda=lambda, mu=mu, t=t3) * # tree 3
  birthdeathprob(a=1, n=0, lambda=lambda, mu=mu, t=t4) * # tree 3
  birthdeathprob(a=1, n=1, lambda=lambda, mu=mu, t=t5) * # tree 3
  6 * # six arrangements
  (1/3) # because the topology is only concordant 1/3 of the time

return(pseudo1discord2)
```

```
}
```
